# Light microscopy based approach for mapping connectivity with molecular specificity

**DOI:** 10.1101/2020.02.24.963538

**Authors:** Fred Y. Shen, Margaret M. Harrington, Logan A. Walker, Hon Pong Jimmy Cheng, Edward S. Boyden, Dawen Cai

**Affiliations:** Medical Scientist Training Program, University of Michigan, Ann Arbor, Michigan, USA; Neuroscience Graduate Program, University of Michigan, Ann Arbor, Michigan, USA; Cellular and Developmental Biology, University of Michigan, Ann Arbor, Michigan, USA; LS&A, Program in Biophysics, University of Michigan, Ann Arbor, Michigan, USA; McGovern Institute, Media Lab, Department of Biological Engineering, and Department of Brain and Cognitive Sciences, Massachusetts Institute of Technology, Cambridge, Massachusetts, USA

## Abstract

Mapping neuroanatomy is a foundational goal towards understanding brain function. Electron microscopy (EM) has been the gold standard for connectivity analysis because nanoscale resolution is necessary to unambiguously resolve chemical and electrical synapses. However, molecular information that specifies cell types is often lost in EM reconstructions. To address this, we devised a light microscopy approach for connectivity analysis of defined cell types called spectral connectomics. We combined multicolor genetic labeling (Brainbow) of neurons with a **m**ult**i-r**ound **i**mmunostaining **Ex**pansion Microscopy (miriEx) strategy to simultaneously interrogate morphology, molecular markers, and connectivity in the same brain section. We applied our multimodal profiling strategy to directly link inhibitory neuron cell types with their network morphologies. Furthermore, we showed that correlative Brainbow and endogenous synaptic machinery immunostaining can be used to define putative synaptic connections between spectrally unique neurons, as well as map putative inhibitory and excitatory inputs. We envision that spectral connectomics can be applied routinely in neurobiology labs to gain insights into normal and pathophysiological neuroanatomy across multiple animals and time points.

## Introduction

Mammalian brains are extraordinarily complex pieces of circuitry, composed of trillions of connections between diverse cell types. Understanding these wiring patterns is a fundamental piece of the puzzle towards understanding how our brains work. A comprehensive wiring diagram of neuronal connections is called a connectome, while the pursuit of a connectome is known as connectomics. Neurons are connected to each other primarily through chemical synapses. They are located where presynaptic and postsynaptic neurons touch, and are composed of structures spanning hundreds of nanometers in size. The presynaptic compartment contains a region that mediates neurotransmitter release called the active zone^1^. The postsynaptic compartment contains a region that is protein dense called the postsynaptic density (PSD), which is comprised of neurotransmitter receptors, scaffolding proteins, and signaling molecules^2^.

Due to their small size, neuroscientists have relied on electron microscopy (EM) to observe chemical synapses and map connections between neurons. EM is a powerful tool that offers unparalleled nanometer resolution^3^, sufficient to observe synaptic structures. Recent works have used EM to define the adult *Drosophila* hemibrain connectome^4^, as well as curate a 92.6 × 94.8 × 61.8 μm^3^ connectome from mouse somatosensory cortex^5^. Despite the recent technological and biological advances with EM, several challenges remain. Molecular information is lost as proteins can only be rarely identified with EM alone. Long imaging timespans generate large datasets that require demanding computational processing and analysis. All of these challenges have placed the use of EM for connectomics beyond the reach of common neurobiology labs. Moreover, because of the aforementioned difficulties in scaling the technology, the use of EM for connectomics is not currently suitable for high throughput experimentation. Given that neuroanatomy can vary between animals and substantially change in disease states, a new approach is needed for connectivity analysis that considers molecular information and is easily scalable.

In contrast to EM, light microscopy (LM) can generate specific molecular information through immunostaining, but lacks the nanoscale resolution to map synapses. Super-resolution light microscopy (srLM) techniques, such as STORM^6–8^ or STED^9^, possess nanoscale resolution and molecular specificity, but are not suitable for thick brain volumes because of slow imaging speeds, photobleaching, and optical distortions. The recent development of expansion microscopy^10–12^, which grants super-resolution to routine LM imaging through isotropic, physical magnification of hydrogel-tissue hybrids, offers an alternative path forward that can combine molecular specificity, nanoscale resolution, and rapid LM in thick brain volumes^13^.

Here, we describe a strategy to obtain high throughput morphology measurements of densely labeled neurons, with integrated molecular and connectivity information for multimodal analysis. We combine a multi-color genetic labeling tool (Brainbow) with a **m**ult**i**-**r**ound **i**mmunostaining **Ex**pansion microscopy (miriEx) strategy to simultaneously profile single neuron morphologies, molecular marker expression, and connectivity in the same brain section. We define the derivation of these properties from hyperspectral fluorescent channels as <u>spectral connectomics</u>, a LM based approach towards mapping neuroanatomy and connectivity with molecular specificity.

## Results

Spectral connectomics is based on the ability to acquire multichannel LM datasets at nanoscale resolution that encode neuron morphologies, cell type profiles, and connectivity. Thus, we needed to develop a strategy for 1) dense labeling of a neuronal population with the ability to unambiguously trace dendrites and axons, 2) multiplexed readout of important protein markers, and 3) multiscale imaging that can range from nanoscale resolution for resolving synapses, to microscale resolution for resolving cell type markers. To do so, we first optimized an expansion microscopy protocol for multi-round immunostaining (miriEx) that let us generate multichannel LM datasets at multiple resolutions. We then added Brainbow to stochastically express fluorescent proteins (FPs) in a cell-type-specific population of neurons^14,15^. The combination of miriEx and Brainbow allowed us to curate hyperspectral, multiscale LM datasets that contain information about molecular markers, neuron morphologies, and synaptic machinery (**Fig 1a**).

**Figure 1.**
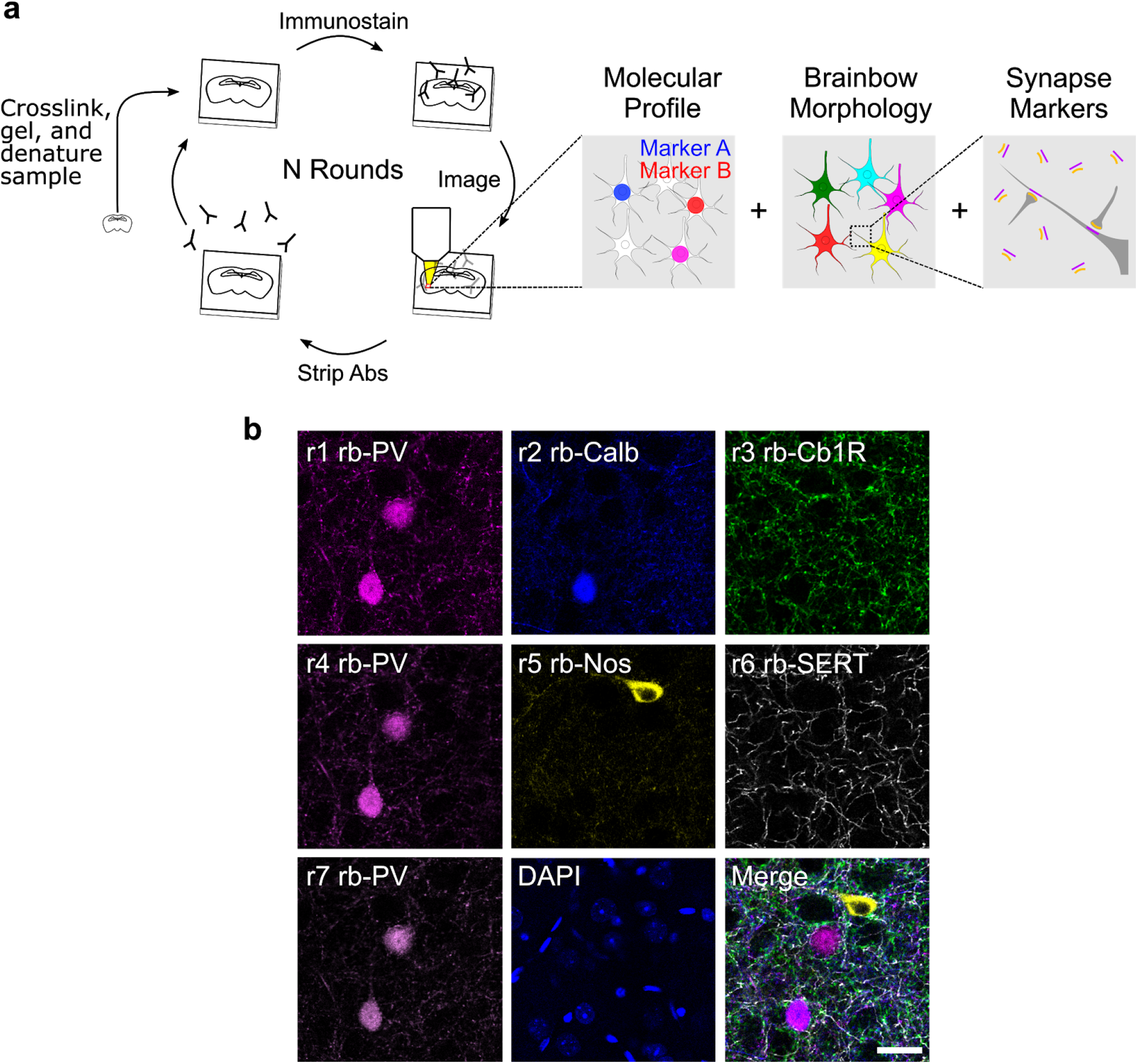
miriEx robustly preserves antigens across multiple rounds of immunostaining, imaging, and stripping. (**a**) miriEx strategy to simultaneously measure molecular profile, morphology, and/or connectivity. Tissue samples are embedded in a hydrogel to create expandable gel-tissue hybrids. The sample can then undergo multiple rounds of immunostaining, imaging, and stripping to measure various neuron properties. We then register and merge the different rounds of imaging to correlate the results. (**b**) 5 distinct rabbit antibodies were used across 7 sequential rounds of immunostaining, with PV re-probed in rounds 4 and 7 to demonstrate retention of antigen. Abs, antibodies; PV, parvalbumin; Calb, calbindin; Cb1R, cannabinoid receptor type 1; Nos, nitric oxide synthase; SERT, serotonin transporter. Scale bar: (**b**) 25 μm (pre-expansion size).

We start by describing the development of miriEx for multiplexed immunostaining. Probing multiple proteins using traditional immunohistochemistry (IHC) is typically limited by host animal species of primary antibodies (most tend to be either mouse or rabbit) and visible light bandwidth, such that detecting more than 4 targets becomes difficult. Recent strategies, such as Immuno-SABER^16^, PRISM^17^, and CODEX^18^ have been developed to overcome these limitations using antibody-DNA barcoding and read out strategies. Alternatively, tissue sections can undergo multiple rounds of routine antibody staining, imaging, stripping, and restaining, as seen in array tomography^19^, CLARITY^20^, MAP^21^, and SHIELD^22^. We adopted the latter strategy and optimized a protein crosslinking protocol to anchor antigens into an expandable hydrogel. Specifically, we used acrylic acid *N*-hydroxysuccinimide ester to modify proteins with acryl groups so they can be crosslinked and polymerized into an expandable hydrogel. We replaced the Proteinase K digestion step from standard expansion microscopy protocols^10,12^ with SDS/heat based denaturation because Proteinase K destroys endogenous proteins. In contrast, SDS/heat treatment preserves endogenous proteins^21^ and is compatible with post-gelation immunostaining. Furthermore, because our denaturation method is similar to that of SDS-PAGE western blots, we found that antibodies already validated for western blots usually work with miriEx (**Table S2**). We also found that the same SDS/heat treatment can efficiently strip antibodies after each round of probing (**Fig. S1**). We validated that gel-tissue hybrids in miriEx expand ~2x in 1x PBS and ~4x in 0.001x PBS respectively (**Fig. S2**).

We then demonstrated that miriEx robustly preserves antigens across multiple rounds of immunostaining and stripping. Conceptually, different rounds of staining can be used to probe different neuronal properties, such as molecular markers, morphology, and/or synaptic markers (**Fig. 1a**). We probed 5 different rabbit antibodies across 7 rounds of imaging in one basolateral amygdala (BLA) sample (**Fig. 1b**), imaging DAPI in each round to use as a fiduciary channel for registration (**Fig. S3**). We found that the same parvalbumin (PV) antibody showed similar staining patterns across multiple rounds with high signal remaining in neurites, indicating that antigen signal can be reliably retrieved, even in later rounds of miriEx. To extend the application of multi-round immunostaining, we showed that miriEx works with formalin-fixed human tissue (**Fig. S4**). We also demonstrated that through multispectral imaging for 3-4 antigens in each round, we could achieve highly multiplexed profiling of 15 different targets in the same piece of mouse striatum tissue (**Fig. S5**).

In the BLA multi-round immunostaining experiment described above, we observed that although the majority of amygdala PV neurons co-expressed calbindin (Calb), some were calbindin negative. Past studies indicate that these PV+/Calb− neurons are axo-axonic cells that specifically innervate the axon initial segment^23^, while PV+/Calb+ neurons represent basket cells that innervate the perisomatic region^24^. Somatostatin (SOM) also marks another broad interneuron subtype in the amygdala, with both Calb positive and negative co-expression^25^. As a result, we chose this system to demonstrate the ability to interface molecular marker information with morphology analysis using miriEx.

To differentiate intermingled neurons *in situ*, we used Brainbow, a technique that relies on the stochastic expression of fluorescent proteins (FPs) to label neighboring neurons in unique colors^14,15^. Importantly, 1) the FPs used are distinct antigens, allowing their signal to be amplified through miriEx immunostaining, 2) the FPs are membrane targeted, which enables better labeling of neuron subcellular morphology^15^, and 3) dense labeling can be achieved to study morphology in a more high throughput manner compared to other techniques that rely on sparse labeling. PV-Cre and SOM-Cre double transgenic animals were generated to allow genetic access to two broad interneuron types. Brainbow AAVs 2/9 were stereotaxically injected in the basolateral amygdala (BLA), and 200 μm sections of tissue were processed with miriEx (**Fig. 2a**). In round 1, three different molecular markers (Calb, PV, and SOM) were probed to define 4 molecular cell types: PV, PV/Calb, SOM, SOM/Calb (**Fig. 2b-e**). In round 2, three Brainbow FPs were immunostained to reveal morphology (**Fig. 2f-i**). DAPI was co-stained as a fiduciary channel for registration of the two rounds. Both rounds of imaging took place with the gel-tissue hybrid expanded ~2x in 1xPBS, giving us an effective imaging resolution of ~150 × 150 × 350 nm^3^. Brainbow AAVs labeled 2 out of 4 (50%) PV neurons, 20 out of 31 (70%) PV/Calb neurons, 7 out of 7 (100%) SOM neurons, and 24 out of 28 (86%) SOM/Calb neurons within a 590 × 404 × 160 μm^3^ volume. Following identification of each Brainbow neuron by its molecular subtype, we reconstructed its dendritic morphology using nTracer^26^, an ImageJ/Fiji plugin for tracing multispectral datasets (**Fig. 2g-h**, **Movie S1**). A total of 53 neurons in the imaged volume was reconstructed across all 4 molecular subtypes and various morphology parameters were analyzed (**Fig. 2l-m**, **Fig. S6**). In agreement with past findings, PV expressing neurons appear to have more complex branching patterns compared with SOM expressing neurons, despite having similar dendritic lengths^25,26^. This type of dense reconstruction enables study of how different neuronal cell types interact with each other anatomically, making it a powerful tool for network morphology analysis.

**Figure 2.**
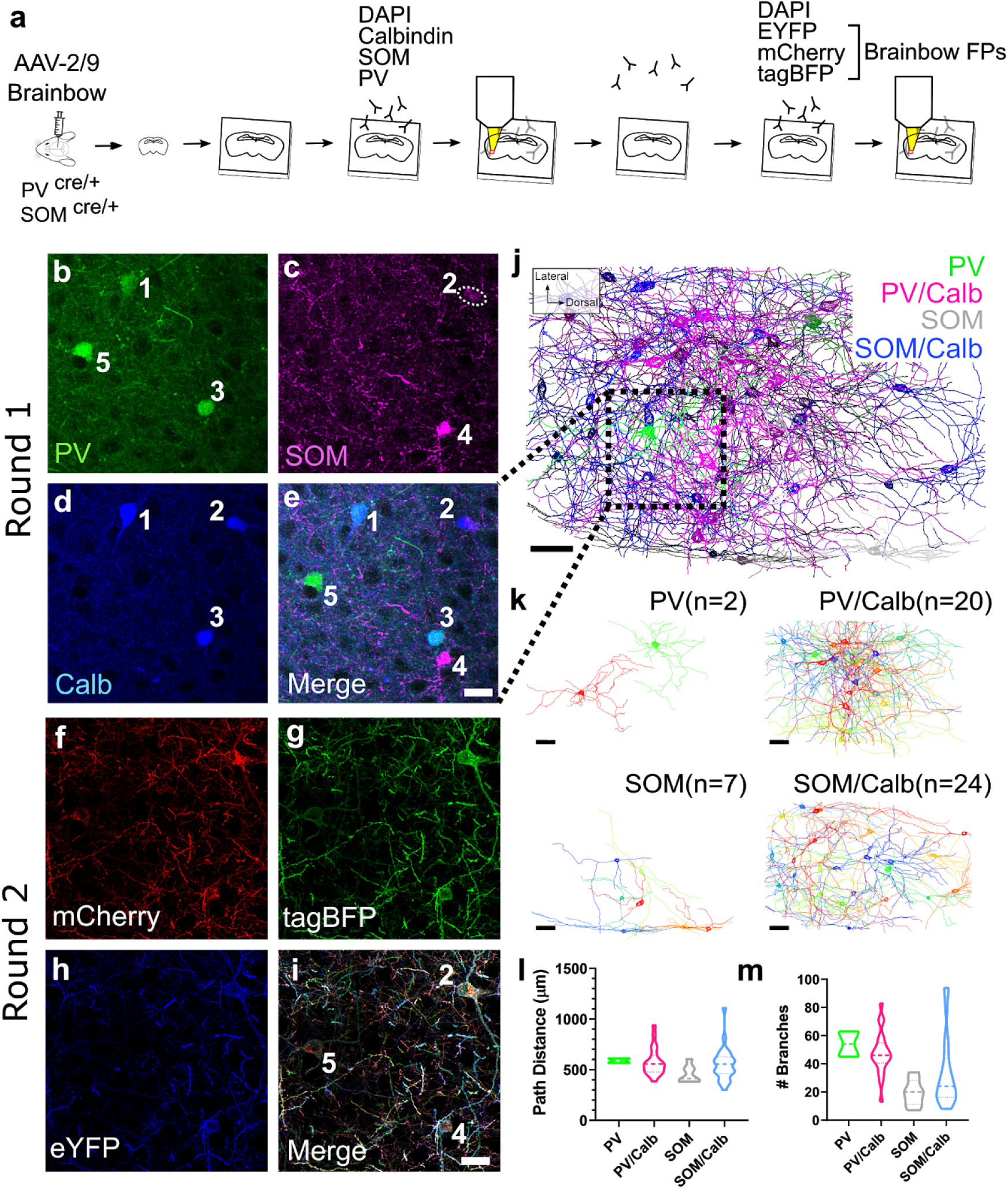
miriEx can be combined with Brainbow to simultaneously profile molecular expression and neuron morphologies. (**a**) Experiment design: Brainbow and molecular markers are imaged across two rounds of immunostaining using DAPI channel for registration. (**b-e**) MIP showing the molecular markers (PV, SOM, Calb) imaged in round 1. Four distinct molecular subtypes could be identified: PV, PV/Calb, SOM, SOM/Calb. (**f-i**) MIP showing the Brainbow channels imaged in round 2. (**j**) nTracer reconstruction of 53 neurons encompassing the 4 subtypes previously identified in a 590 × 404 × 160 μm^3^ volume. The red square represents the field of view seen in **b-i**. (**k**) Individual nTracer reconstructions based on molecular subtype. (**l**) Total path distance plotted for each of the molecular subtypes. (**m**) Number of dendritic branches plotted for each of the molecular subtypes. MIP, maximum intensity projection; PV, parvalbumin; Calb, calbindin; SOM, somatostatin. Scale bars: (**e,i**) 25 μm (pre-expansion size). (**j,k**) 50 μm (pre-expansion size).

While understanding the projection patterns of axons and dendrites is an important aspect of mapping neuroanatomy, another central objective is to understand how neurons connect. Chemical synapses are hundreds of nanometers in size and serve as bridges of communication between neurons^7^. Measuring synapses using conventional light microscopy (LM) techniques is difficult as the distance between synaptic structures and neuronal boundaries can be smaller than the diffraction limit^6^. Recently, expansion microscopy has been shown to be a viable strategy for resolving synaptic structures and assigning them to neurons^13,27,28^. Consequently, we combined Brainbow with miriEx to measure synaptic structures located at the junctions between different neurons in an effort to define connectivity using LM.

We first confirmed that miriEx was compatible with endogenous synaptic machinery immunostaining by probing for Gephyrin (inhibitory PSD), Homer1 (excitatory PSD), and Bassoon (presynaptic active zone) in layer 4 somatosensory cortex (**Fig. S7**). ~4x expansion of the sample gave us an effective imaging resolution of ~70 × 70 × 200 nm^3^ using confocal microscopy. We found inhibitory Gephyrin-Bassoon synapse pairs to be less common (21%) than excitatory Homer1-Bassoon synapse pairs (77%). Importantly, only ~2% of the Bassoon puncta were not paired with Gephyrin or Homer1, which were mutually exclusive (**Fig. S7**). This gave us confidence that the Gephyrin and Homer1 antibodies we used mark inhibitory and excitatory synapses almost completely. We also showed that over 90% of inhibitory and excitatory synapses are greater than 300 nm away from their closest neighbor, allowing us to reliably distinguish neighboring synapses at our imaging resolution (**Fig. S7**).

We then packaged Brainbow in AAV-PHP.eB^29^ serotype to efficiently transduce neurons systemically across the brain via intravenous injection. We retro-orbitally injected AAV-PHP.eB Brainbow in PV-Cre mice and found that we had near complete coverage of PV neurons in somatosensory cortex (**Fig. S8**). 100% of the Brainbow labeled neurons were positive for PV immunostaining indicating that our labeling strategy was both highly sensitive and specific (**Fig. S8**). 100 μm sections of somatosensory cortex were processed with miriEx. In round 1, three Brainbow FPs were stained and the sample was expanded and layer 4 was imaged (**Fig. 3a-b**). In round 2, a presynaptic marker (Bassoon), inhibitory postsynaptic marker (Gephyrin), and EYFP were stained, and the sample was again expanded and imaged in layer 4 (**Fig. 3c**). We observed that Bassoon-Gephyrin pairs could be resolved and were located at axosomatic and axodendritic contact points between Brainbow labeled PV neurons (**Fig. 3d-e**, **Movie S2**). Measuring the line profile of these putative synapses revealed that Gephyrin, the post-synaptic Brainbow membrane, and Bassoon were arranged in the expected order (**Fig. 3h-i**). The distance between Gephyrin and Bassoon puncta was between 100-200 nm and matched previous reports^7,27^. The synaptic cleft between pre- and post-synaptic Brainbow membranes is tens of nanometers wide and could not be resolved at this resolution, but the trio of Gephyrin-Brainbow-Bassoon signal still gave us confidence to define putative inhibitory synapses between Brainbow labeled PV neurons. Moreover, our approach towards spectral connectomics allowed us to eliminate false positives that would not be possible with diffraction-limited LM. Neuronal contacts have previously been used as correlates for synaptic sites ^26,30^, but we show an example of a close apposition lacking synaptic machinery that would be falsely classified as a connection (**Fig. 3g, k**). Another type of error that could arise is the wrongful assignment of a physically close axon that merely passes through and doesn’t form a synapse (**Fig. 3f, j**). Given that axons can be small caliber with bouton sizes that are hundreds of nanometers^31^, diffraction limited LM may confuse neighboring axons, leading to synapse assignment mistakes.

**Figure 3.**
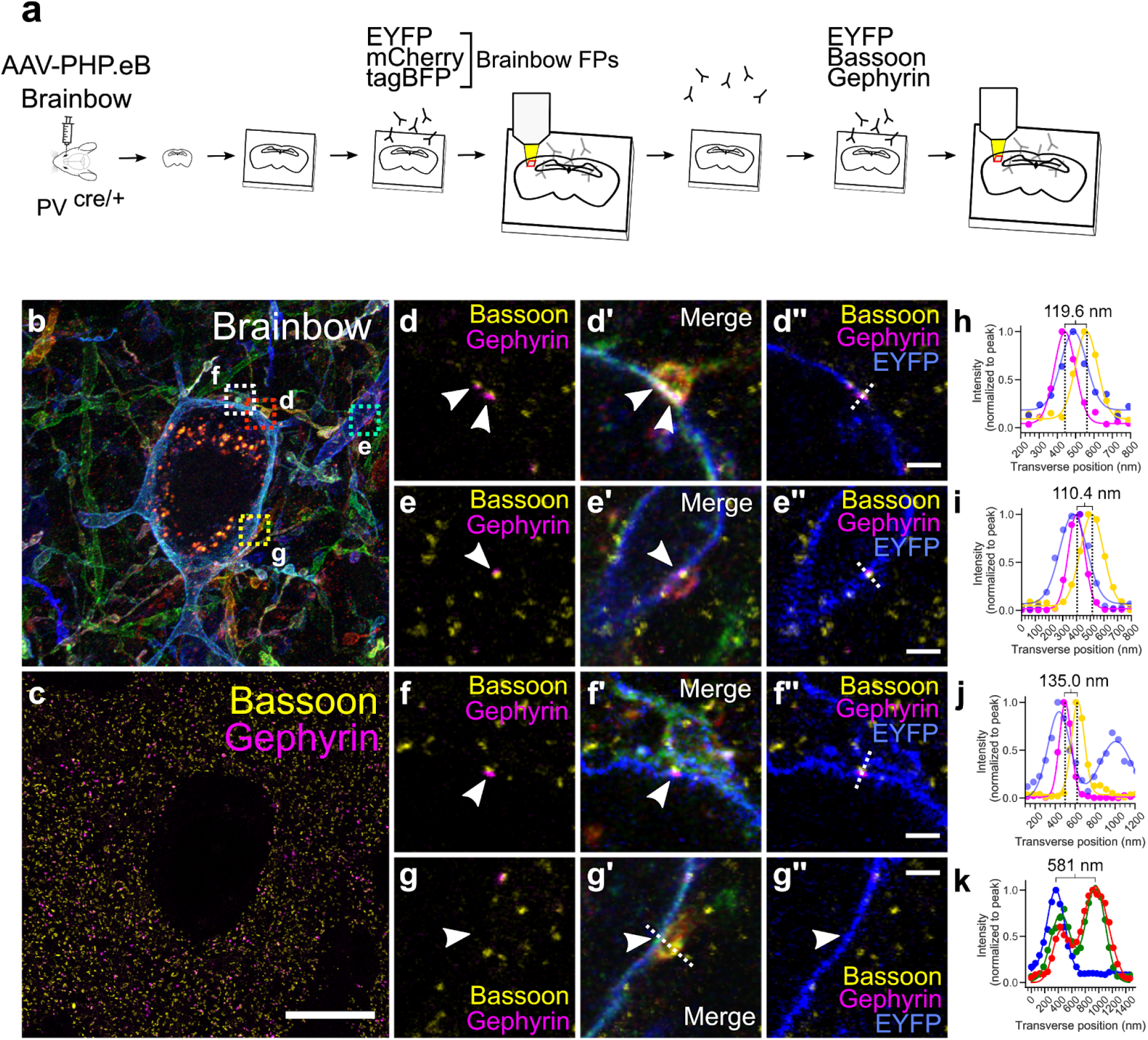
Putative synaptic connections can be defined between Brainbow neurons. (**a**) Experimental design: Brainbow FPs and endogenous synaptic markers are imaged across two rounds of immunostaining using the EYFP channel for registration. (**b**) MIP showing the Brainbow channels imaged in round 1. (**c**) MIP showing the Bassoon and Gephyrin puncta imaged in round 2. (**d-d’’**) Single slice zoomed inset of red square in **b**. Example of two axo-somatic synapses where the orange axon contacts the blue soma. Two Gephyrin-Bassoon pairs (white arrows) are shown within the axonal bouton. (**e-e’’**) Single slice zoomed inset of cyan square in **b**. Example of an axo-dendritic synapse where the red axon contacts the blue dendrite. White arrow points to the Gephyrin-Bassoon pair. (**f-f’’**) Single slice zoomed inset of white square shown in **b**. Example of an axon that is physically close, but doesn’t actually actually form a synapse. White arrow points to the Gephyrin-Bassoon pair. (**g-g’’**) Single slice zoomed inset of yellow square in **b**. Example of an axo-somatic contact that lacks Gephyrin-Bassoon machinery (white arrow) and is not a true synapse. (**h-k**) Normalized line intensity profiles of dotted lines drawn in **d-g** respectively. Distances between Gephyrin and Bassoon peaks or between two Brainbow cell membranes were measured and shown respectively for **h-j** or **k**. Scale bars: (**c**) 10 μm (pre-expansion size). (**d-g**) 1 μm (pre-expansion size).

Following the first two rounds of miriEx that probed Brainbow and synaptic markers, a third round of miriEx was performed to probe SOM, adding information about molecular cell types to the dataset (**Fig. S9**). By doing so, we show the ability to map PV connectivity onto a variety of cell types. However, we note that it remains difficult to accurately assign synapses without both the presynaptic and postsynaptic Brainbow membranes. Without a postsynaptic SOM neuron membrane label, it is challenging to tell if the PV axon synapses directly on the soma or on an unlabeled, small caliber dendrite sandwiched in between. As a result, we did not attempt to analyze PV to SOM connectivity.

After validating that putative inhibitory synapses could be identified between Brainbow labeled PV neurons, we set out to trace the axons and dendrites of 8 PV neurons whose somas were located inside a ~100 × 100 × 60 μm^3^ imaging volume (**Fig. 4a**). We then traced every Brainbow labeled PV axon that innervated these 8 PV neurons, and annotated all the putative inhibitory synapses (Gephyrin-Brainbow-Bassoon trios) that we could identify. 189 axons were traced and 422 molecularly specific (PV-PV) putative synapses were defined (**Fig. 4b**, **Movie S3**). First we analyzed the connections between these 8 PV neurons by plotting their connectivity matrix (**Fig. 4c**). We observed that neuron 37 innervated 4 other PV neurons, matching previous reports of local PV-PV connectivity^32^. Interestingly, while local PV-PV connection is common in our dataset, we did not find local reciprocal inhibition between two PV neurons. Next we added 189 traced PV axons, whose somas were not located in this volume, to the same connectivity matrix (**Fig. 4d**). We found multiple examples of presynaptic PV axons that innervate more than one postsynaptic PV neuron. However, many of the dimly labeled PV axons could only be reliably traced for a short distance, leaving the full extent of their synaptic connections uncovered. This likely skewed the connectivity matrix towards under-representing PV axons that synapse onto multiple postsynaptic PV neurons. Compared to traditional monochromic labeling, Brainbow labeling provides spectral information to identify the source of innervating axons. For example, we can be confident that two spectrally unique axons come from different PV neurons even without tracing them back to their somas. Same colored axons are more difficult to interpret as they could come from different neurons that happen to have similar colors, or they could originate from the same neuron branch *outside* our imaging volume. Nevertheless, we can still use color to estimate the upper and lower bounds when asking the convergence question of *how many unique PV neurons innervate one PV neuron*. The upper bound is determined by the number of innervating axons, while the lower bound is determined by the number of unique colors these axons possess (**Fig. 4e**). To determine the number of unique colors, we plotted their RGB color on a ternary plot and used an elbow plot to conservatively estimate the number of k-means color clusters (**Fig. S10**). Within our dataset volume, the average upper and lower bound of unique PV neurons converging onto one PV neuron are estimated as 27.5 ± 12.6 and 10.9 ± 4.5 respectively (n = 8, mean ± standard deviation).

**Figure 4.**
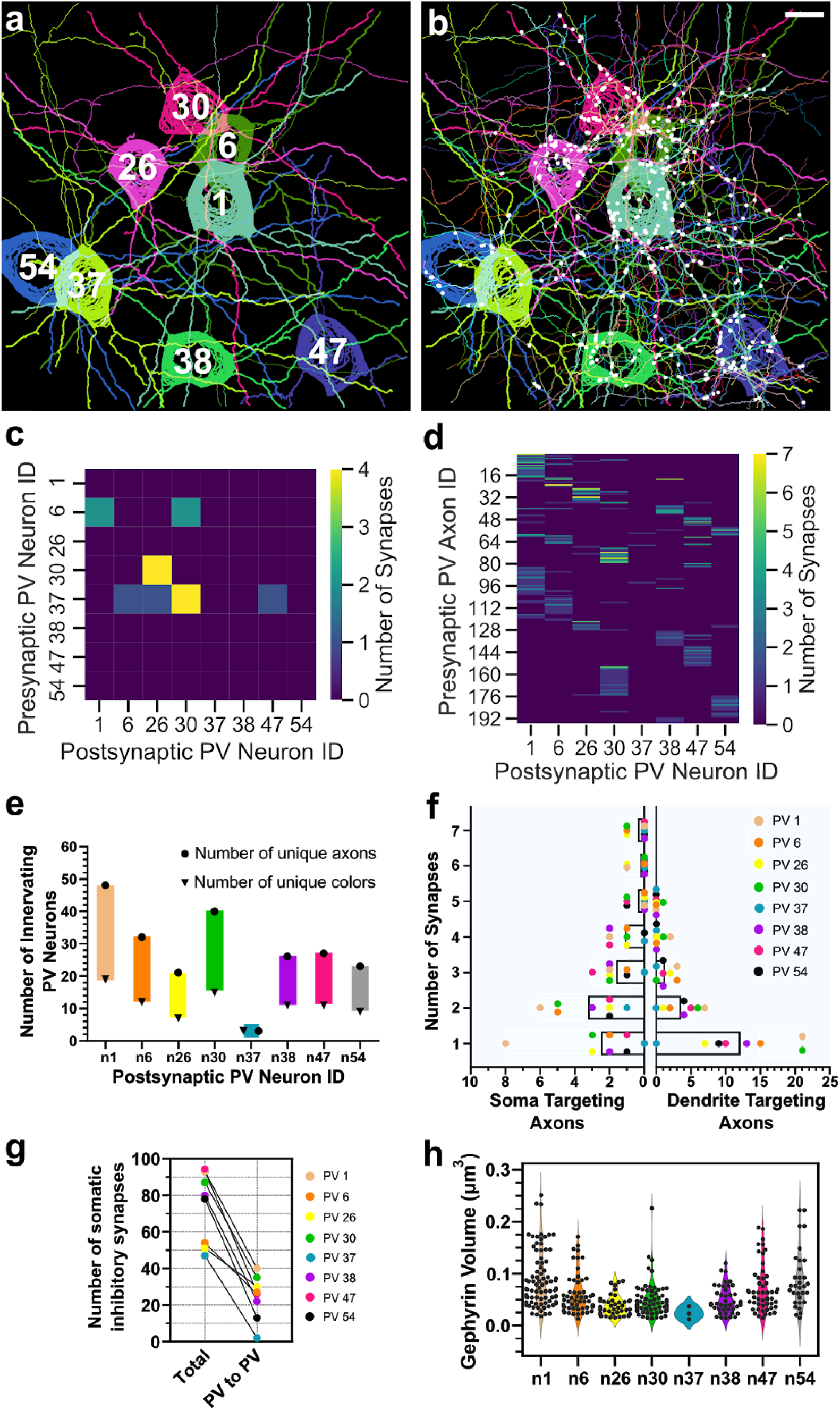
Molecularly specific analyses can be performed on spectral connectomics data. (**a**) MIP of 8 Brainbow labeled PV neurons reconstructed with nTracer. Thick neurites represent dendrites; thin neurites represent axons. The neuron IDs are overlaid. (**b**) Same as **a**, plus all of the other 189 innervating axons overlaid and 422 identified synapses marked with white circles. (**c**) Connectivity matrix between the 8 reconstructed PV neurons. (**d**) Connectivity matrix as in **c**, plus the 189 innervating axons. (**e**) Plot of the maximum and minimum number of unique presynaptic PV neurons that innervate each of the 8 postsynaptic PV neurons. Circles represent the number of spatially distinct PV axons that formed synapses, while triangles represent the number of unique colors that could be identified from the innervating axons. (**f**) Distribution of the number of soma targeting and dendrite targeting axons as a function of the number of synapses they form. Each of the 8 afferent PV neurons are color coded and plotted with the average represented by the bar graph. (**g**) Total number of somatic inhibitory synapses were plotted relative to the PV to PV subset for each of the 8 postsynaptic PV neurons. (**h**) Inhibitory PSD (Gephyrin) size plotted for all the PV to PV synapses found on each of the 8 postsynaptic PV neurons. Scale bars: 10 μm (pre-expansion size).

Next we shifted our attention from the distribution of axons to the distribution of synapses. We analyzed the distribution of PV-PV inhibitory synapses by splitting them into two spatial compartments: somatic and dendritic (**Fig. 4f**). We plotted the number of soma or dendrite targeting axons based on the number of putative synapses they provide. Cortical PV neurons are known as basket cells and tend to innervate the perisomatic region of other neurons^33^. While we confirmed the existence of putative PV-PV axosomatic synapses, a larger number was actually axodendritic. The soma targeting axons most commonly formed two putative synapses, but in contrast, the majority of the dendrite targeting axons formed one putative synapse. **Fig. 3d** shows an example of a single axonal bouton providing two putative axosomatic synapses. We then counted the total number of putative inhibitory synapses on the soma and plotted it alongside those that were annotated as PV-PV (**Fig. 4g**). We observed that approximately 33.3 ± 16.5 % of the somatic inhibitory synapses were PV-PV (n = 8, mean ± standard deviation). Because inhibitory synapses are plastic and can dynamically remodel^33–35^, we wanted to determine if there were any differences in synapse size, a known correlate for synapse strength^36,37^, between the 8 postsynaptic PV neurons. We looked at the distribution of Gephyrin volume for all PV-PV synapses for each of the 8 postsynaptic PV neurons and found they were largely consistent with no major differences (**Fig. 4h**). Next, we analyzed individual postsynaptic PV neurons and asked whether axons that provide more synapses are correlated with increased PSD sizes. Again, we observed PSD sizes were consistent and invariant of the number of synapses an axon formed (**Fig. S11**).

Previous studies performing *in situ* synapse measurements tend to focus on excitatory neurons because dendritic spines are commonly used as a proxy for excitatory input^38^. Changes in spine size and density are often used as a correlate to structural synaptic plasticity^38^. Studies of inhibitory neuron synapses are more challenging and underrepresented since many of them are aspiny with no obvious morphological correlate. To meet this challenge, we simultaneously labeled aspiny BLA PV neurons with Brainbow, excitatory (Homer1), and inhibitory (Gephyrin) PSD markers (**Fig. 5a**). In round 1, three Brainbow FPs were immunostained and the sample was expanded and imaged (**Fig. 5b**). In round 2, Homer1 and Gephyrin were immunostained along with EYFP as a fiduciary channel (**Fig. 5c**). Expansion of the sample gave us the resolution to optically resolve individual synaptic puncta along the Brainbow membrane (**Fig. 5d-i**). We reconstructed the dendritic morphology of 5 PV cells in a 220 × 220 × 85 μm^3^ imaging volume, and annotated all the excitatory (915 ± 274, mean ± standard deviation, n=5) and inhibitory (409 ± 141, mean ± standard deviation, n=5) PSDs to create a synaptic input map for each cell (**Fig. 5j-k, Movie S4**). We propose that these synaptic input maps may be another useful metric to distinguish cell types. For example, we intriguingly observed that three out of five PV neurons possess a skewed distribution of excitatory vs. inhibitory inputs, while the other two PV neurons possess a more balanced distribution (**Fig. 5l**). The excitatory vs. inhibitory input ratio fundamentally influences a neuron’s role in the circuit, and is another measure of connectivity that can be used to distinguish neuronal subtypes.

**Figure 5.**
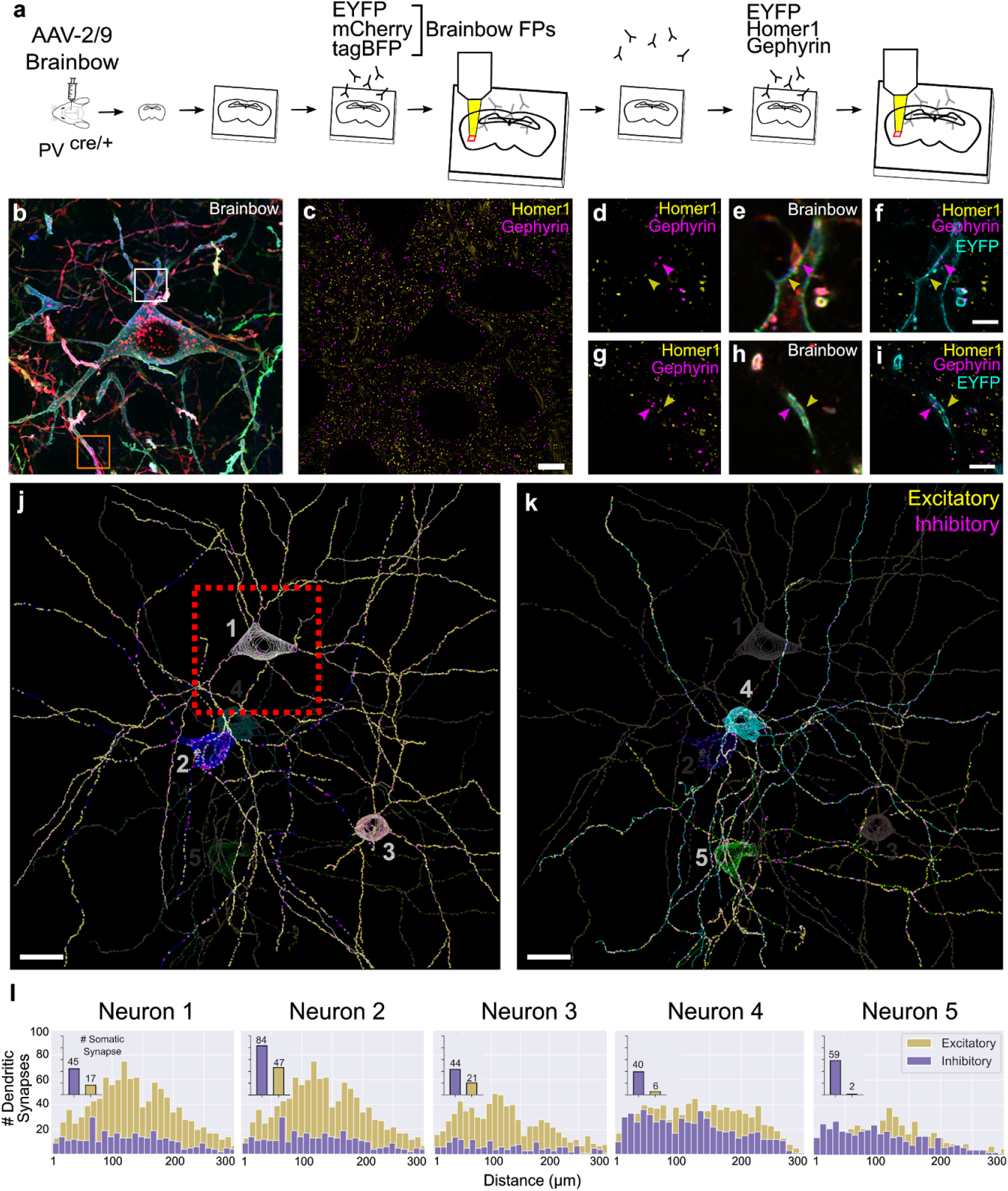
Putative excitatory and inhibitory synapses of aspiny inhibitory neurons can be quantified. (**a**) Experiment design: Brainbow and endogenous PSD markers are imaged across two rounds of immunostaining using the EYFP channel for registration. (**b**) MIP of the three Brainbow channels imaged in round 1. (**c**) Single slice of excitatory (Homer1) and inhibitory (Gephyrin) PSDs imaged in round 2. (**d-i**) Zoomed in synaptic marker + Brainbow single slice images of the white square (top row) and orange square (bottom row) respectively. Yellow and purple arrows point to excitatory and inhibitory synaptic puncta respectively. (**j-k**) nTracer reconstruction of 5 neurons’ morphology and putative synaptic inputs. The red box represents the field of view seen in **b** and **c**. (**l**) Histogram of the number of putative dendritic synapses as a function of dendritic length away from the soma for each neuron. The inset bar graph represents the number of putative somatic synapses for each neuron. Scale bars: (**c**) 10 μm (pre-expansion size), (**f,i**) 2.5 μm (pre-expansion size), (**j,k**) 20 μm (pre-expansion size).

## Discussion

In summary, we present a LM based approach for connectivity analysis with molecular specificity, termed spectral connectomics, developed by combining Brainbow and miriEx technologies. We demonstrated multimodal measurements of morphology, molecular markers, and connectivity in a single mammalian brain section. We showed that miriEx supports robust preservation of antigens to enable multiple immunostaining, imaging, and stripping cycles. This allowed us to reconstruct the morphology and specify the molecular subtype of 53 different BLA interneurons in the same brain section. Adding endogenous immunostaining of pre and postsynaptic pairs enabled us to define putative synaptic connections between molecularly specified neurons. We traced 8 postsynaptic PV neurons,189 innervating PV axons, and 422 PV-PV inhibitory synapses, and performed connectomic analyses similar to previous EM studies. Finally, we showed that Brainbow could be combined with endogenous PSD immunostaining to quantify inhibitory and excitatory synaptic input onto individual aspiny interneurons.

miriEx enhances the potential of using Brainbow for mapping neuroanatomy. Depending on the labeling density, diffraction-limited LM may be insufficient for tracing small caliber Brainbow neurites^8,26^. Expansion microscopy allowed us to resolve these neurites effectively for morphology and connectivity analysis. The membrane targeted FPs also proved to be an useful landmark when probing synaptic machinery. Both pre and postsynaptic membranes were required along with immunostaining for pre and postsynaptic machinery to identify putative synapses. In addition, membrane targeted FPs are shown to be better than cytosolic FPs, which may not diffuse as well, for depicting small subcellular structures^25^. Two challenges that remain with combining Brainbow and miriEx are the limited color diversity and dilution of membrane FP signal. Throughout our experiments, we immunostained for only three out of four possible FPs. We can already observe tens of easily distinguishable color barcodes, which can be exponentially increased in the future by probing more FPs. For example, 5 FPs were used in the recently developed Bitbow, a “digital” format of Brainbow^39^. Expansion of the sample is important for increasing imaging resolution, but comes at the cost of diluting antibody signal. For instance, ~4x expansion results in a ~64 fold reduction of antibody signal. As a result, axons can become dim and challenging to trace long distances. That said, we expect a more photon efficient volumetric imaging modality, such as light sheet microscopy, will yield better signal to noise under the same labeling conditions than confocal microscopy used in our study. Moving forward, it will be important to optimize FP expression along axonal membranes, as well as explore alternative signal amplification technologies, such as immuno-SABER^16^.

A critical component of our spectral connectomics strategy is molecular specificity, which comes from two sources: endogenous protein immunostaining and choice of Cre-driver line. The ability to probe endogenous molecular markers with miriEx increases the flexibility of studying multiple neuronal cell types simultaneously. For example, future work can focus on using a broad transgenic line (e.g. GAD2-Cre) that can be molecularly specified by additional immunostaining (PV, SOM, VIP, etc). Furthermore, miriEx immunostaining can be extended to investigate the molecular content of synapses. Bassoon, Gephyrin, and Homer1 are only a subset of the synaptic machinery that can be probed, which includes neurotransmitter transporters, receptors, gap junctions, and ion channels. The rich library of Cre-driver lines can give us a genetic handle on studying specific cell types. Labeling a subset from the global population of neurons also helps to constrain and focus our connectivity analysis. For example, we only quantified and analyzed 422 PV-PV synapses out of ~60,000 inhibitory synapses in our dataset (~0.7% of the total). Because we asked molecularly specific, targeted questions of PV-PV connectivity, we reduced the burden of tracing neurons and annotating synapses to a scale that was manageable by manual curation (~40 human work hours).

Moreover, many Cre-driver lines are actually composed of different neuronal subtypes^40^. Our definitions of cell types are constantly changing with the introduction of new technologies. We have progressed from using unimodal criteria (*e.g*. morphology, physiology, or molecular expression) to using a combination of these properties in a multimodal fashion^41^. Recent studies have demonstrated the power of this multimodal approach to curate and refine definitions of inhibitory and excitatory neuronal cell types in visual cortex^42,43^. Similarly, our multimodal strategy for measuring morphology, molecular markers, and connectivity can be also applied towards refining cell type definitions across the brain. More specifically, integrating connectivity information will enhance our understanding of how input/output properties inform cell type definitions. In addition, spectral connectomics is compatible with techniques that measure physiological properties, such as patch clamp electrophysiology and *in vivo* calcium recordings, for more comprehensive analysis.

Finally, we envision spectral connectomics to supplement EM as an alternative strategy for mapping connectivity that is accessible to neurobiology labs. miriEx and Brainbow use commercially available, “off the shelf”, reagents, and LM is a prevalent imaging modality across many labs. Data sizes are an order of magnitude smaller compared to EM datasets despite containing multiple channels (~20 GB vs ~200 GB for a similar sized volume^5^). Targeted analysis makes it possible to map connectivity in a reasonable amount of time, and application of automated reconstruction and synapse segmentation methods will further simplify this process. EM based connectomics currently remains the gold standard for generating complete reference connectomes of animal brains, but we hope that spectral connectomics can be employed for validating molecularly specific circuit motifs and testing hypotheses of how neuroanatomy dynamically changes with perturbations. We imagine that in the future, spectral connectomics can be scaled up to generate comprehensive connectomes. This likely will come from developing new Brainbow designs to globally label neurons^29^, whole brain expansion microscopy protocols, whole brain LM methods^44,45^, whole brain immunolabeling techniques^46,47^, and new computational pipelines for automated neuron reconstruction^48^ and synapse segmentation^49^. Although challenging, many of these puzzle pieces are already being developed by the community. When combined in the spectral connectomics framework, they can fit together to reveal a mammalian connectome.

## Methods

### Mouse Lines

All experiments were carried out in compliance with the National Institutes of Health Guide for the Care and Use of Laboratory Animals. Our study protocol was reviewed and approved by the University of Michigan Institutional Animal Care & Use Committee. The transgenic mice used in this study were: PV-Cre (Jackson stock no. 008069), SOM-Cre (Jackson stock no. 013044), VGAT-Cre (Jackson stock no. 016962), and Thy1-YFP-H (Jackson stock no. 003782). P28 to P56 male and female mice were used for virus injections.

### Brainbow AAV injections

Brainbow3.0 AAV-2/9 and AAV-PhP.EB were obtained from Addgene and University of Michigan vector core, respectively. Transgenic mice (PV-Cre/SOM-Cre or PV-Cre) were anesthetized continuously with isofluorane and mounted on a stereotaxic frame. TagBFP-EYFP and mCherry-TFP virus were mixed together to reach a concentration of 1E12 gc/mL individually, of which 500 nL was injected at 100 nL/min using a capillary pipette backfilled with mineral oil at +3.5 ML and −1.7 AP relative to bregma, 2.7 DV from pia surface. Afterward, the pipette was left in place for 5 min. for the virus to diffuse, before slowly retracting out of the brain. We waited 3-4 weeks for virus expression before perfusing the animals. Brainbow3.0 AAV-PhP.EB was used for retro-orbital injection to systematically label neurons throughout the brain. 50 uL of mixed virus (1E12 gc total each for TagBFP-EYFP and mCherry-TFP) was injected into the retro-orbital sinus. We waited 3-4 weeks for virus expression before perfusing the animals.

### Mouse perfusion and tissue sectioning

Mice were anesthetized with tribromoethanol and perfused transcardially with ice-cold 1xPBS, followed by 4% PFA in 1xPBS. The brains were dissected and postfixed in 4% PFA in 1xPBS overnight shaking at 4 °C. The next day, brians were washed in 1xPBS before slicing on a vibratome (Leica VT1000s) at 100 μm or 200 μm thickness.

### miriEx protocol

Acrylic acid N-hydroxysuccinimide ester (AAx, Sigma A8060) was prepared by dissolving in N,N-Dimethylformamide to 125 mM. Tissue samples were incubated with 1-5 mM Aax in a MBS buffer (100 mM MES, 150 mM NaCl, pH 6) with 0.1% Triton X-100 shaking overnight at 4 °C. The next day, the samples were washed 3x with 1xTBS (Bio-rad 1706435) for 1 hour each to quench the reaction. The samples were then incubated in monomer solution (5.3% Sodium Acrylate, 4% Acrylamide, 0.1% Bis-Acrylamide, 0.5% VA-044, in 1x PBS) shaking overnight at 4 °C. The next day, the samples were gelled for 2.5 hours at 37 °C in a humidity chamber by cover slipping the sample surrounded by monomer solution in a gelling chamber. The gel-tissue hybrids were then carefully cut out and denatured at 70 °C overnight in denaturing buffer (200mM SDS dissolved in 1xTBS). The next day, denaturing buffer was washed out by incubating the samples 4x for 2 hours each in 0.1% PBST (1x PBS with 0.1% Triton X-100) shaking at 50 °C.

### Immunohistochemistry of regular tissue sections

For immunostaining of non-gelled samples, tissue sections were first blocked and permeabilized in StartingBlock (Thermo 37538) with 1% Triton X-100 shaking overnight at 4 °C. Tissues were washed the next day 3x with 1xPBS for 1 hour each. Tissues were then incubated with primary antibodies diluted in 0.5% PBST for 3 days at 4 °C. After washing 3x with 0.5% PBST for 1 hour each at RT, tissue sections were then incubated with secondary antibody diluted in 0.5% PBST for 2 days at 4 °C. Tissues were then mounted with Vectashield (Vectorlabs H-1000) and imaged. Primary and secondary antibody choice and concentrations can be found in **Table S1**.

### Immunohistochemistry of gelled tissue sections

Blocking and permeabilization of gel-tissue hybrids were skipped as they were found to minimally decrease background. Gel-tissue samples were incubated with primary antibody diluted in 0.1% PBST w/ 0.02% azide (PBSTz) at either RT or 37 °C. They were washed 3x with 0.1% PBST w/ 0.02% Azide at RT for 1 hour each, and then incubated with secondary antibody diluted in 0.1% PBSTz at either RT or 37°C. Primary and secondary antibody choice and concentrations can be found in **Table S1**. Vendor information can be found in **Table S2 and S3**.

### Gel expansion and fluorescence microscopy

miriEx gel-tissue hybrids were incubated in 0.001x PBS three times for 45 min. each for saturated expansion to ~4x. They were then mounted in Poly-L-Lysine coated 6 cm dishes (Corning 354517) and submerged in 0.001x PBS. All confocal LM was performed using an upright Zeiss LSM780. Water immersion objectives (10X/NA0.4 or 20x/NA1.0) were directly lowered into the PBS solution over the sample for imaging. More imaging details are presented in **Table S1**.

### Antibody elution

miriEx gel-tissue hybrids that previously have undergone immunostaining, expansion, and imaging were shrunken by washing in 1xPBS 3x for 1 hour each. They were then put in denaturing buffer (200 mM SDS dissolved in 1xTBS) at 70 °C overnight. Samples were washed the next day 4x for 2 hours each in 0.1% PBST shaking at 50 °C before undergoing the next round of immunostaining.

### Human brain sample processing with miriEx

Formalin-fixed human brain samples were obtained from the University of Michigan Brain Bank. A 1 cm slab of sensory cortex was macro dissected and washed in 1x PBS at 4 °C overnight. The sample was then sectioned into 100 μm slices before processing with miriEx.

### Image preprocessing

Specific image processing steps for each imaging dataset are listed in **Table S1**, and are briefly explained below.

Stitching of multi-tile datasets was performed using the BigStitcher^41^ ImageJ/Fiji plugin. The dataset was loaded and converted to HDF5 format, and the tiles were arranged in the order they were imaged with 10% overlap. The Stitching Wizard was used to calculate pairwise shifts using phase correlation, verify links, and undergo global optimization. Affine refinement was then performed with a high threshold. The resulting interest points were used for non-rigid refinement during advanced image fusion.

Chromatic aberration was corrected using the Detection of Molecules (DoM) ImageJ/Fiji plugin. To calibrate the DoM plugin, 0.5 μm TetraSpeck fluorescent beads were mounted on a slide and imaged on a Zeiss LSM780 confocal microscope. More specifically, we calibrated the orange channel (540-600 nm) excited with a 543 laser and far-red channel (630-700 nm) excited with a 633 laser to the green channel (480-540 nm) excited with a 488 laser. Calibration was performed for both the 10x and 20x objectives.

Histogram matching was done to normalize intensity between z-slices in image stacks using the nTracer Align-Master ImageJ/Fiji plugin. A high signal to noise z-slice (usually at the top or bottom of the stack) was chosen as the reference slice for which the rest of the stack was normalized to. All three Brainbow FP channels were histogram matched to the same reference slice.

### Image registration and alignment between rounds

Following data preprocessing as described above, the fiducial marker channels from different rounds were loaded into ImageJ/Fiji Big Warp^50^ plugin for rough, initial alignment. After this, the fiducial marker channels were registered through a B-spline transformation using Elastix^51,52^. The resulting transformation was applied to each individual channel to create a merged image hyperstack.

### Morphology reconstruction, synapse identification, and analysis

nTracer, an ImageJ/Fiji plugin, was used to trace somas, dendrites, and axons of Brainbow labeled neurons. The morphology reconstructions were exported in SWC format, and a custom Python script was used to render skeleton visualizations in TIFF format from SWC files. Putative synapses were identified and manually marked using ROI manager in ImageJ/Fiji. The list of X, Y, Z synapse coordinates was saved to a CSV file and linked with their parent neurons by adding an additional data column in the SWC file marking synaptic locations along the dendrite or axon. Blender 2.81 (Blender Foundation; www.blender.org) or 3Dscript^53^ was used to generate movies. Morphology features (i.e. number of stems, bifurcations, branches, etc.) were calculated by importing SWC files into Vaa3D’s Global Neuron Feature plugin^54^. Sholl analysis was performed using ImageJ/Fiji Sholl Analysis plugin. Automatic segmentation of Gephyrin volumes was achieved through a custom Python script by first applying an Otsu thresholding step, followed by watershed segmentation of Gephyrin puncta. Plots were made in GraphPad Prism 8 or through Matplotlib Python scripts.

### miriEx expansion measurement

100 μm Thy1-YFP-H sections were mounted in Vectashield and imaged using confocal microscopy. The sections were then washed in 1xPBS to remove the vectashield and were processed with miriEx. An anti-GFP antibody was used to label native YFP signal. After miriEx, the samples were imaged in 1xPBS in the same location as before. They were then further expanded in 0.001x PBS (3x washes for 45 min. each) and imaged again. Elastix^51^ was used to calculate an affine transformation between different expansion states, and the X, Y, and Z scaling factors were averaged to measure the expansion factor.

## Data Availability

The data that support the findings of this study are available from the corresponding author upon reasonable request.

## Code Availability

Custom code for analysis of image processing and data analysis is available from the corresponding author upon reasonable request.

## Acknowledgements

We thank Nigel Michki and Douglas Roossien for providing comments, editing, and discussion of the manuscript. We thank Harsimranjit Sekhon for insightful discussion. We thank Andy Lieberman for providing formalin-fixed human brain tissue. We thank Bo Duan and Lorraine Horwitz for providing SOM-Cre mice and genotyping assistance. FYS acknowledges support by National Institutes of Health (NIH) 1F31NS11184701. DC acknowledges support by NIH 1UF1NS107659, 1R01MH110932, and 1RF1MH120005, and National Science Foundation NeuroNex-1707316. ESB acknowledges support by Lisa Yang, John Doerr, the Open Philanthropy Project, and Schmidt Futures, and NIH 1R01NS102727, 1R01EB024261, and 1R01MH110932.

## Author contribution

FYS and DC conceived the project, designed the experiments, and wrote the manuscript with input from all authors. FYS performed and analyzed all miriEx and Brainbow experiments. HPJC and FYS traced the 53 BLA PV and SOM neurons. LAW wrote code for automatic segmentation of Gephyrin puncta and helped create the BLA molecular subtypes video. MMH and FYS quantified Gephyrin-Bassoon and Homer1-Bassoon pairs, and PV and Brainbow overlap. ESB supported ExM development. DC initiated and supervised the project.

## References

1. Südhof, T. C. The Presynaptic Active Zone. Neuron 75, 11–25 (2012).

2. Sheng, M. & Kim, E. The Postsynaptic Organization of Synapses. Cold Spring Harb. Perspect. Biol. 3, (2011).

3. Xu, C. S., Pang, S., Hayworth, K. J. & Hess, H. F. Enabling FIB-SEM Systems for Large Volume Connectomics and Cell Biology. bioRxiv 852863 (2019) doi:10.1101/852863.

4. Xu, C. S. et al. A Connectome of the Adult Drosophila Central Brain. bioRxiv 2020.01.21.911859v1 (2020) doi:10.1101/2020.01.21.911859v1.

5. Motta, A. et al. Dense connectomic reconstruction in layer 4 of the somatosensory cortex. Science 366, (2019).

6. Sigal, Y. M., Speer, C. M., Babcock, H. P. & Zhuang, X. Mapping Synaptic Input Fields of Neurons with Super-Resolution Imaging. Cell 163, 493–505 (2015).

7. Dani, A., Huang, B., Bergan, J., Dulac, C. & Zhuang, X. Super-resolution Imaging of Chemical Synapses in the Brain. Neuron 68, 843–856 (2010).

8. Rust, M. J., Bates, M. & Zhuang, X. Sub-diffraction-limit imaging by stochastic optical reconstruction microscopy (STORM). Nat. Methods 3, 793–796 (2006).

9. Hell, S. W. & Wichmann, J. Breaking the diffraction resolution limit by stimulated emission: stimulated-emission-depletion fluorescence microscopy. Opt. Lett. 19, 780–782 (1994).

10. Tillberg, P. W. et al. Protein-retention expansion microscopy of cells and tissues labeled using standard fluorescent proteins and antibodies. Nat. Biotechnol. 34, 987–992 (2016).

11. Chen, F., Tillberg, P. W. & Boyden, E. S. Expansion microscopy. Science 347, 543–548 (2015).

12. Chozinski, T. J. et al. Expansion microscopy with conventional antibodies and fluorescent proteins. Nat. Methods 13, 485–488 (2016).

13. Gao, R. et al. Cortical column and whole-brain imaging with molecular contrast and nanoscale resolution. Science 363, eaau8302 (2019).

14. Livet, J. et al. Transgenic strategies for combinatorial expression of fluorescent proteins in the nervous system. Nature 450, 56–62 (2007).

15. Cai, D., Cohen, K. B., Luo, T., Lichtman, J. W. & Sanes, J. R. Improved tools for the Brainbow toolbox. Nat. Methods 10, 540–547 (2013).

16. Saka, S. K. et al. Immuno-SABER enables highly multiplexed and amplified protein imaging in tissues. Nat. Biotechnol. 37, 1080–1090 (2019).

17. Guo, S.-M. et al. Multiplexed and high-throughput neuronal fluorescence imaging with diffusible probes. Nat. Commun. 10, 4377 (2019).

18. Goltsev, Y. et al. Deep Profiling of Mouse Splenic Architecture with CODEX Multiplexed Imaging. Cell 174, 968–981.e15 (2018).

19. Micheva, K. D. & Smith, S. J. Array Tomography: A New Tool for Imaging the Molecular Architecture and Ultrastructure of Neural Circuits. Neuron 55, 25–36 (2007).

20. Chung, K. et al. Structural and molecular interrogation of intact biological systems. Nature 497, 332–337 (2013).

21. Ku, T. et al. Multiplexed and scalable super-resolution imaging of three-dimensional protein localization in size-adjustable tissues. Nat. Biotechnol. 34, 973–981 (2016).

22. Park, Y.-G. et al. Protection of tissue physicochemical properties using polyfunctional crosslinkers. Nat. Biotechnol. 37, 73–83 (2019).

23. Vereczki, V. K. et al. Synaptic Organization of Perisomatic GABAergic Inputs onto the Principal Cells of the Mouse Basolateral Amygdala. Front. Neuroanat. 10, (2016).

24. McDonald, A. J. & Betette, R. L. Parvalbumin-containing neurons in the rat basolateral amygdala: morphology and co-localization of Calbindin-D28k. Neuroscience 102, 413–425 (2001).

25. McDonald, A. J. & Mascagni, F. Immunohistochemical characterization of somatostatin containing interneurons in the rat basolateral amygdala. Brain Res. 943, 237–244 (2002).

26. Roossien, D. H. et al. Multispectral tracing in densely labeled mouse brain with nTracer. Bioinformatics 35, 3544–3546 (2019).

27. Chang, J.-B. et al. Iterative expansion microscopy. Nat. Methods 14, 593–599 (2017).

28. Lee, K.-S., Vandemark, K., Mezey, D., Shultz, N. & Fitzpatrick, D. Functional Synaptic Architecture of Callosal Inputs in Mouse Primary Visual Cortex. Neuron 101, 421–428.e5 (2019).

29. Chan, K. Y. et al. Engineered AAVs for efficient noninvasive gene delivery to the central and peripheral nervous systems. Nat. Neurosci. 20, 1172–1179 (2017).

30. Holler-Rickauer, S., Köstinger, G., Martin, K. A. C., Schuhknecht, G. F. P. & Stratford, K. J. Structure and function of a neocortical synapse. bioRxiv 2019.12.13.875971 (2019) doi:10.1101/2019.12.13.875971.

31. Chéreau, R., Saraceno, G. E., Angibaud, J., Cattaert, D. & Nägerl, U. V. Superresolution imaging reveals activity-dependent plasticity of axon morphology linked to changes in action potential conduction velocity. Proc. Natl. Acad. Sci. 114, 1401–1406 (2017).

32. Jiang, X. et al. Principles of connectivity among morphologically defined cell types in adult neocortex. Science 350, (2015).

33. Villa, K. L. et al. Inhibitory Synapses Are Repeatedly Assembled and Removed at Persistent Sites In Vivo. Neuron 90, 662–664 (2016).

34. Flores, C. E. & Méndez, P. Shaping inhibition: activity dependent structural plasticity of GABAergic synapses. Front. Cell. Neurosci. 8, (2014).

35. Crosby, K. C. et al. Nanoscale Subsynaptic Domains Underlie the Organization of the Inhibitory Synapse. Cell Rep. 26, 3284–3297.e3 (2019).

36. Nusser, Z., Hájos, N., Somogyi, P. & Mody, I. Increased number of synaptic GABA(A) receptors underlies potentiation at hippocampal inhibitory synapses. Nature 395, 172–177 (1998).

37. Lim, R., Alvarez, F. J. & Walmsley, B. Quantal size is correlated with receptor cluster area at glycinergic synapses in the rat brainstem. J. Physiol. 516, 505–512 (1999).

38. Yuste, R. & Bonhoeffer, T. Morphological Changes in Dendritic Spines Associated with Long-Term Synaptic Plasticity. Annu. Rev. Neurosci. 24, 1071–1089 (2001).

39. Veling, M. W. et al. Identification of Neuronal Lineages in the Drosophila Peripheral Nervous System with a “Digital” Multi-spectral Lineage Tracing System. Cell Rep. 29, 3303–3312.e3 (2019).

40. Tasic, B. et al. Shared and distinct transcriptomic cell types across neocortical areas. Nature 563, 72–78 (2018).

41. Yuste, R. et al. A community-based transcriptomics classification and nomenclature of neocortical cell types. ArXiv 190903083 Q-Bio (2019).

42. Gouwens, N. W. et al. Classification of electrophysiological and morphological neuron types in the mouse visual cortex. Nat. Neurosci. 22, 1182–1195 (2019).

43. Gouwens, N. W. et al. Toward an integrated classification of neuronal cell types: morphoelectric and transcriptomic characterization of individual GABAergic cortical neurons. bioRxiv 2020.02.03.932244 (2020) doi:10.1101/2020.02.03.932244.

44. Li, A. et al. Micro-Optical Sectioning Tomography to Obtain a High-Resolution Atlas of the Mouse Brain. Science 330, 1404–1408 (2010).

45. Hillman, E. M. C., Voleti, V., Li, W. & Yu, H. Light-Sheet Microscopy in Neuroscience. Annu. Rev. Neurosci. 42, 295–313 (2019).

46. Renier, N. et al. iDISCO: A Simple, Rapid Method to Immunolabel Large Tissue Samples for Volume Imaging. Cell 159, 896–910 (2014).

47. Yun, D. H. et al. Ultrafast immunostaining of organ-scale tissues for scalable proteomic phenotyping. bioRxiv 660373 (2019) doi:10.1101/660373.

48. Januszewski, M. et al. High-precision automated reconstruction of neurons with flood-filling networks. Nat. Methods 15, 605–610 (2018).

49. Huang, G. B., Scheffer, L. K. & Plaza, S. M. Fully-Automatic Synapse Prediction and Validation on a Large Data Set. Front. Neural Circuits 12, (2018).

50. Bogovic, J. A., Hanslovsky, P., Wong, A. & Saalfeld, S. Robust registration of calcium images by learned contrast synthesis. in 2016 IEEE 13th International Symposium on Biomedical Imaging (ISBI) 1123–1126 (2016). doi:10.1109/ISBI.2016.7493463.

51. Klein, S., Staring, M., Murphy, K., Viergever, M. A. & Pluim, J. P. W. elastix: A Toolbox for Intensity-Based Medical Image Registration. IEEE Trans. Med. Imaging 29, 196–205 (2010).

52. Shamonin, D. P. et al. Fast Parallel Image Registration on CPU and GPU for Diagnostic Classification of Alzheimer’s Disease. Front. Neuroinformatics 7, (2014).

53. Schmid, B. et al. 3Dscript: animating 3D/4D microscopy data using a natural-language-based syntax. Nat. Methods 16, 278–280 (2019).

54. Peng, H., Ruan, Z., Long, F., Simpson, J. H. & Myers, E. W. V3D enables real-time 3D visualization and quantitative analysis of large-scale biological image data sets. Nat. Biotechnol. 28, 348–353 (2010).

