## Supplemental Figures for "Light microscopy based approach for mapping connectivity with molecular specificity"

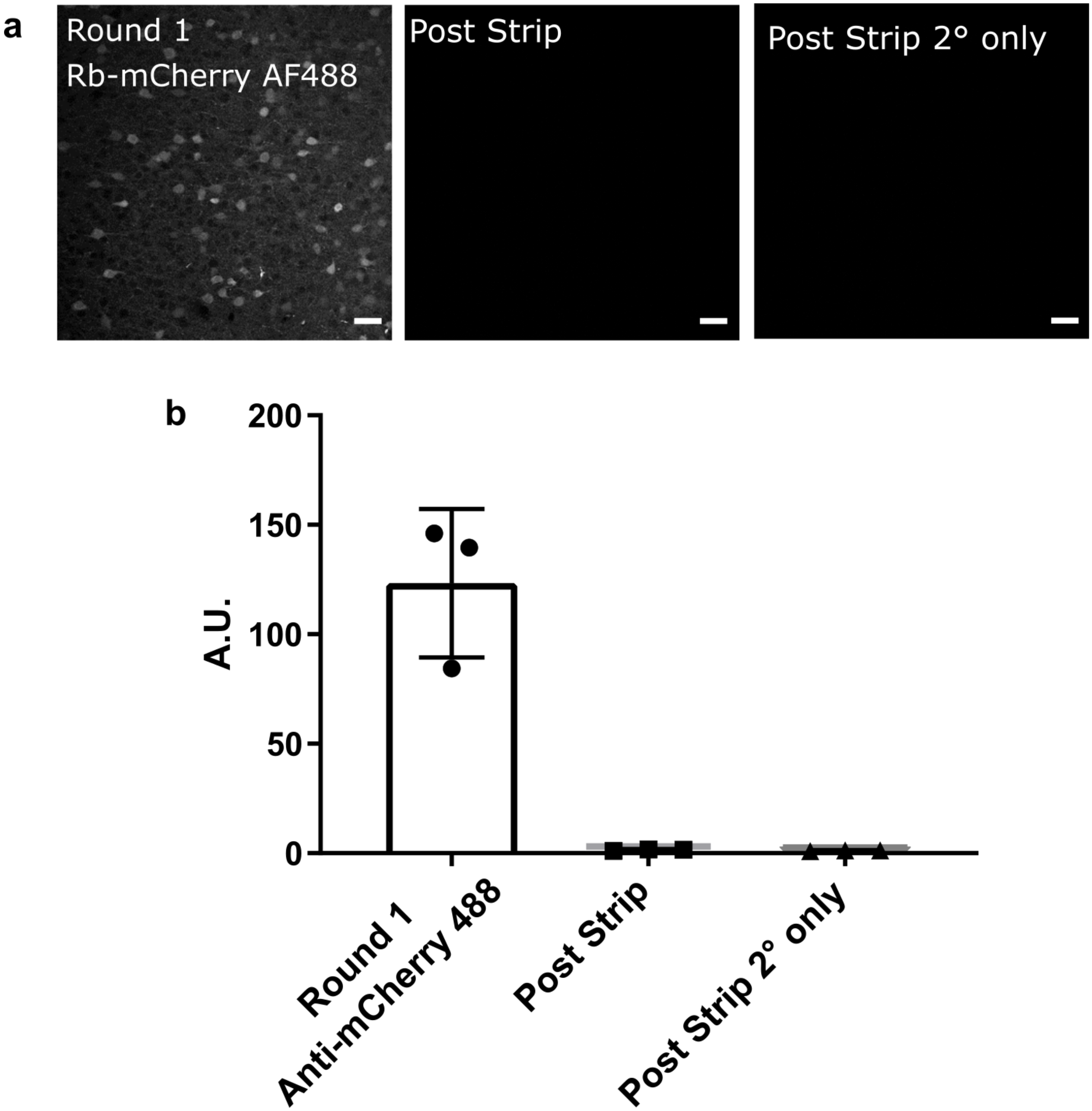


**Supplemental Fig 1. Heat and SDS are efficient at eluting antibody from miriEx gel-tissue hybrids.** (**a**) VGAT-Cre x Ai14 tissue sections were processed with miriEx and TdTomato+ neurons were immunostained with rb-mCherry antibody. The samples were then stripped using heat/SDS and re-imaged. The samples were then stained with secondary antibody only to check that the primary antibody was stripped off. (**b**) Quantification of fluorescence intensity seen in **a** (n=3 samples). Scale bars: (**a**) 50 μm (pre-expansion size).

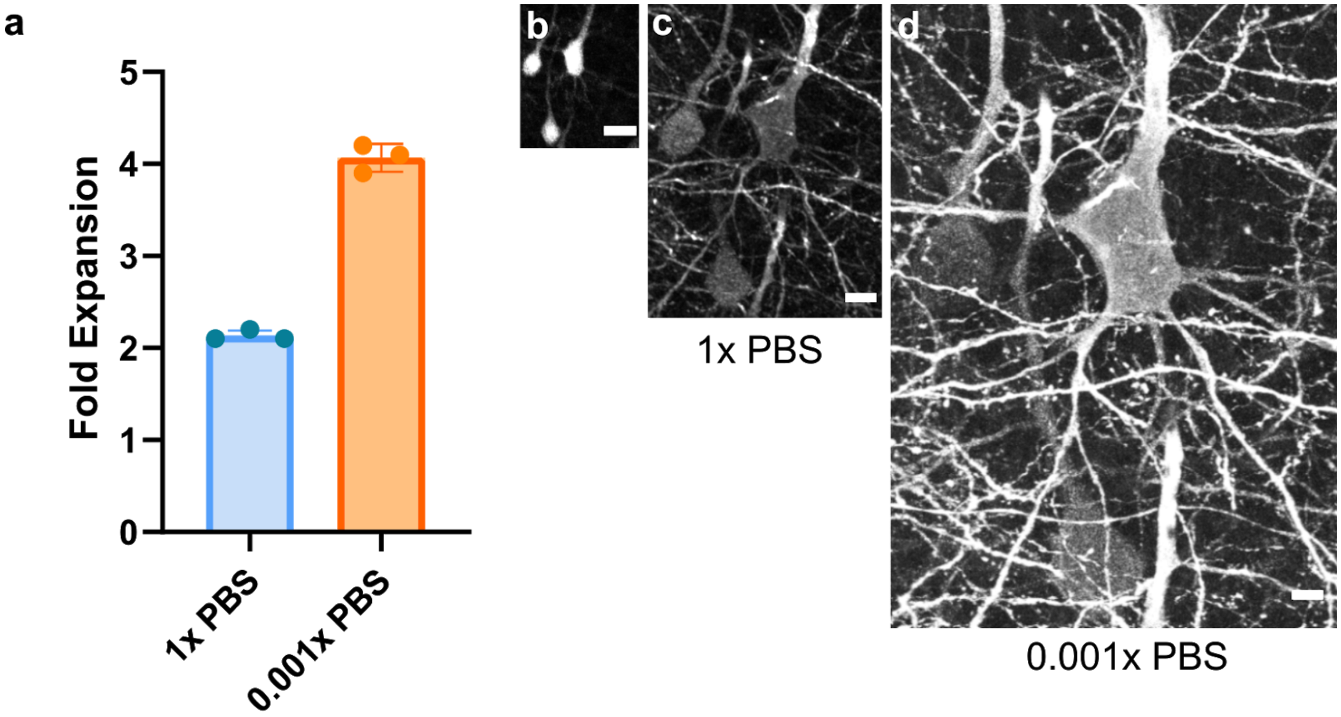


**Supplemental Fig 2. miriEx gel-tissue hybrid expansion.** (**a**) miriEx gels expand approximately ~2 fold in 1xPBS, and ~4 fold in 0.001xPBS (n=3 samples). (**b-d**) Same field of view imaged in three conditions: before miriEx processing, after miriEx processing in 1x PBS, and after miriEx processing in 0.001x PBS respectively. Scale bars: (**b-d**) 20um.

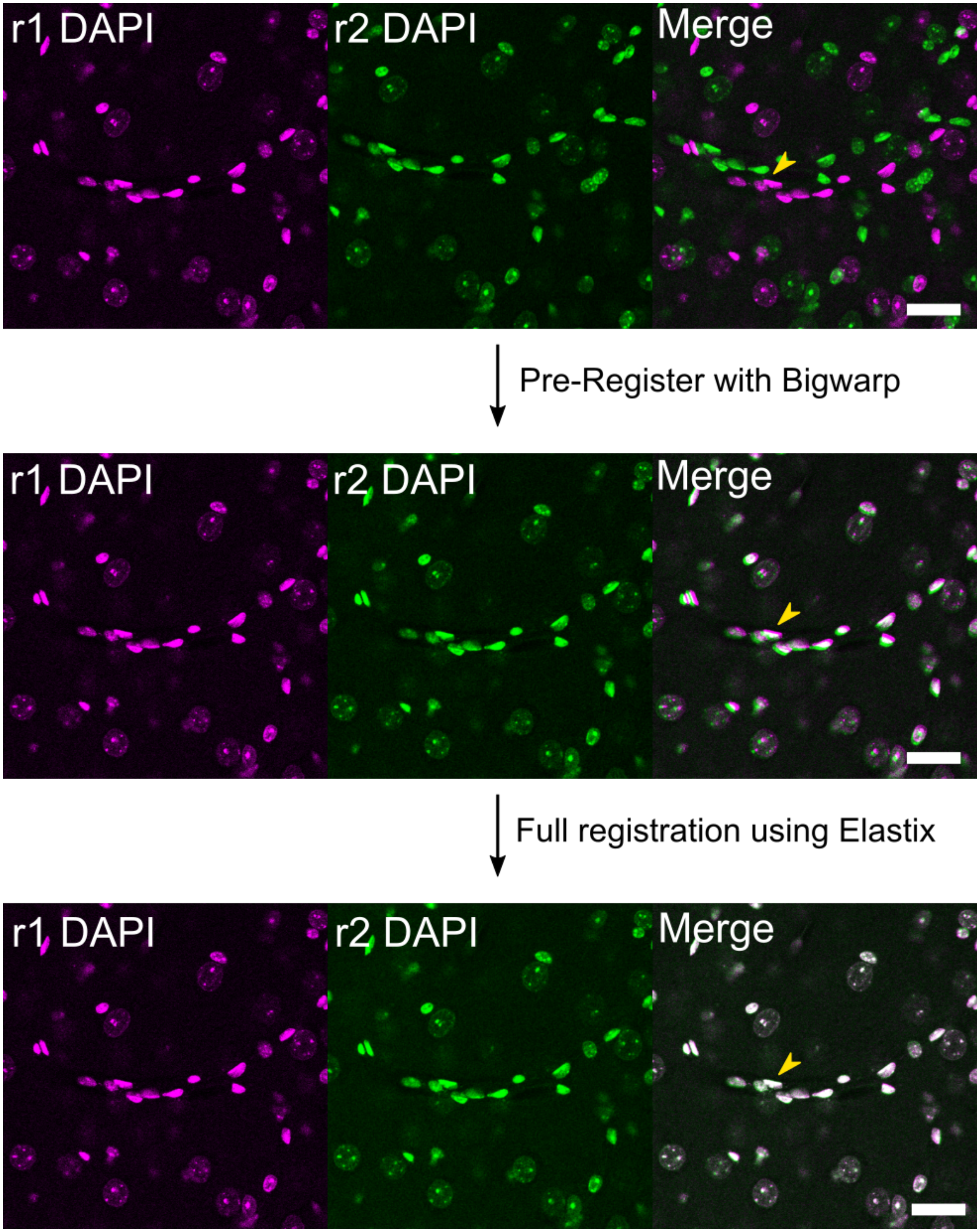


**Supplemental Fig 3. Image registration pipeline between different rounds of miriEx.** Scale bars: 25 µm (pre-expansion size).

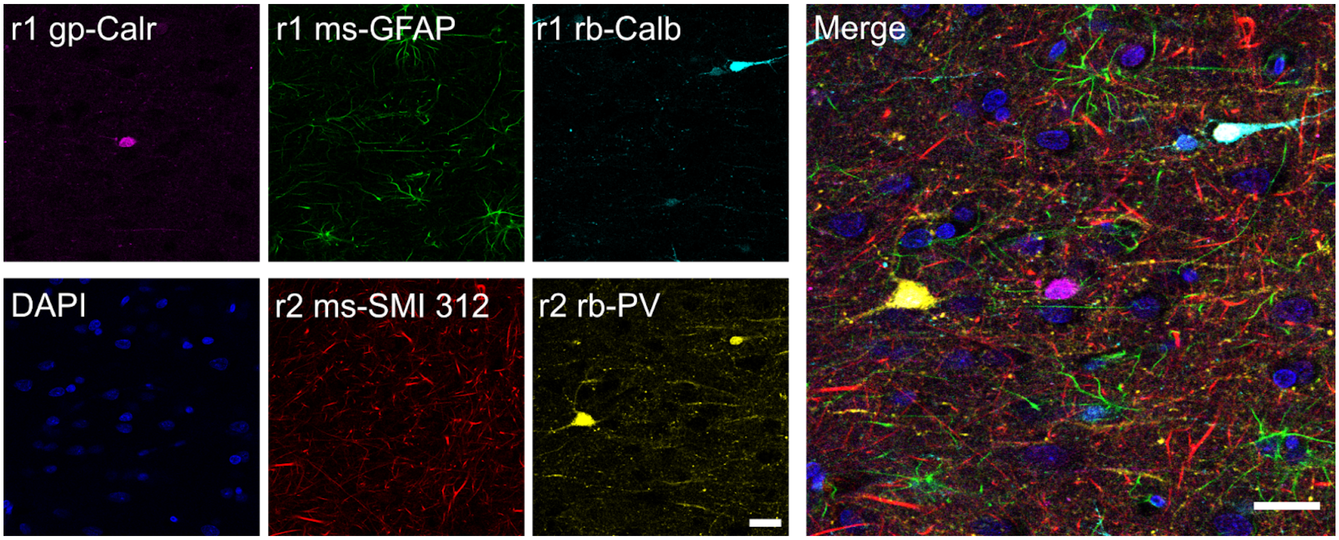


**Supplemental Fig 4. miriEx enables multiplexed immunostaining in formalin fixed human brain tissue.** Calr, Calretinin; PV, parvalbumin; Calb, calbindin; GFAP, glial fibrillary acidic protein. Scale bars: 25 μm (pre-expansion size).

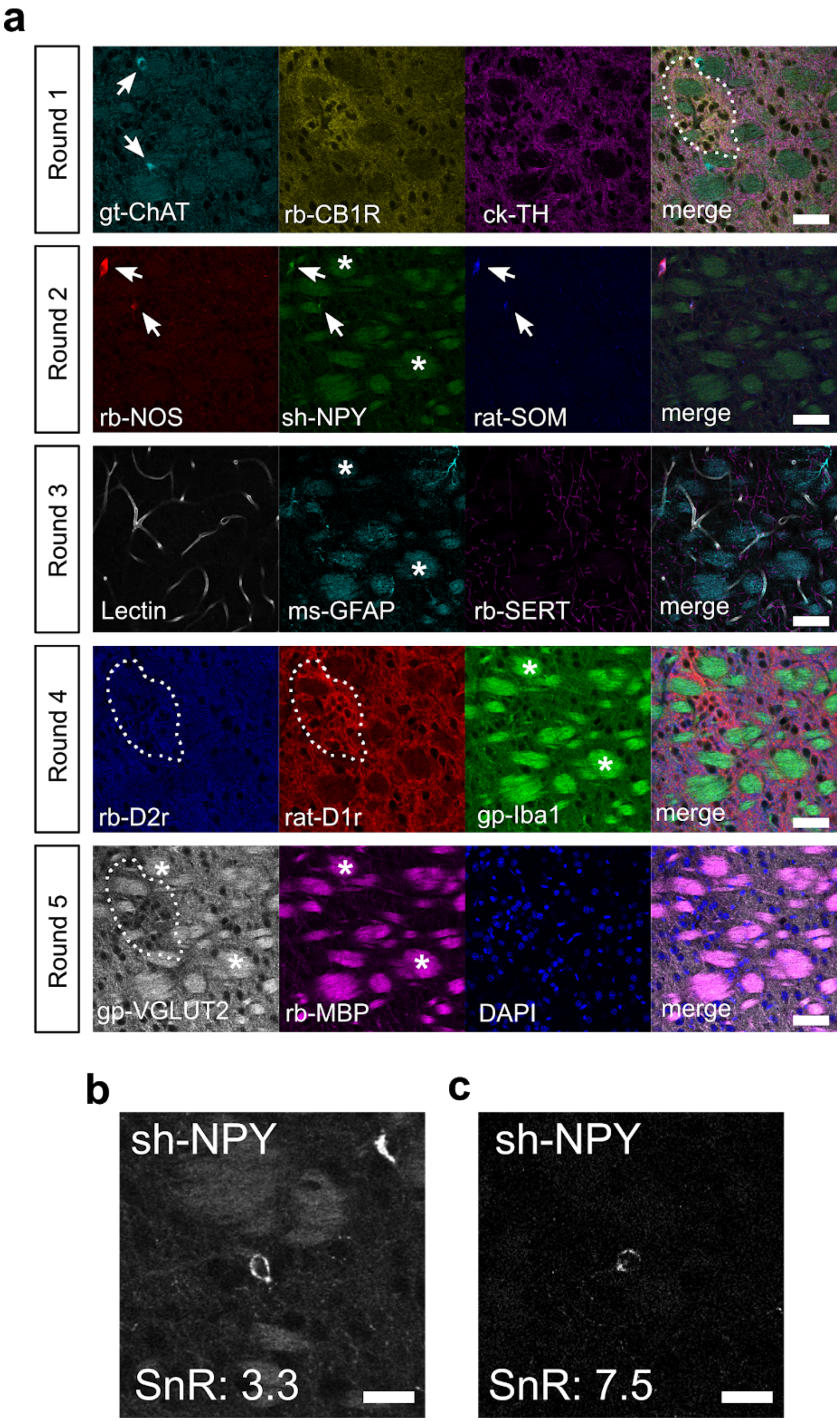
a
Round 1
gt-ChAT
rb-CB1R
ck-TH
merge
Round 2
rb-NOS
sh-NPY
rat-SOM
merge
Round 3
Lectin
ms-GFAP
rb-SERT
merge
Round 4
rb-D2r
rat-D1r
gp-Iba1
merge
Round 5
gp-VGLUT2
rb-MBP
DAPI
merge
b
sh-NPY
SnR: 3.3
c
sh-NPY
SnR: 7.5

**Supplemental Fig 5. miriEx enables highly multiplexed molecular profiling.** (**a**) A total of 15 different antigens across 5 rounds were probed and imaged with confocal microscopy. DAPI was stained in each round to use as a fiduciary channel for registration. CB1R expression marks patches, a histochemically distinct compartment within the striatum (dashed white outline, round 1). NOS, NPY, and SOM expression largely overlap (round 2). D1r expression is higher in striatal patches, while D2r and VGLUT2 expression is higher in the surrounding matrix compartment (round 4). White arrows point to cell bodies with positive molecular marker expression. Asterisks point to myelin bundles immunostained by myelin binding protein (MBP, round 5), typically seen in coronal striatal sections. However, myelin bundles can also be non-specifically stained by secondary antibodies as high background in other rounds. (**b**) An example of NPY immunostaining with high background in striatum. (**c**) Demonstration that lowering the secondary antibody concentration two fold from 1:500 to 1:1000 decreases background and increases signal to noise (SnR). ChAT, choline acetyltransferase; CB1R, cannabinoid receptor type 1; TH, tyrosine hydroxylase; NOS, nitric oxide synthase; NPY, neuropeptide Y; SOM, somatostatin; GFAP, glial fibrillary acidic protein; SERT, serotonin transporter; D1r, dopamine receptor D1; D2r, dopamine receptor D2; Iba1, ionized calcium binding adaptor molecule 1; VGLUT2, vesicular glutamate transporter 2; MBP, myelin basic protein;  Scale bars: (**a**) 50 µm (pre-expansion size). (**b-c**) 25 µm (pre-expansion size).

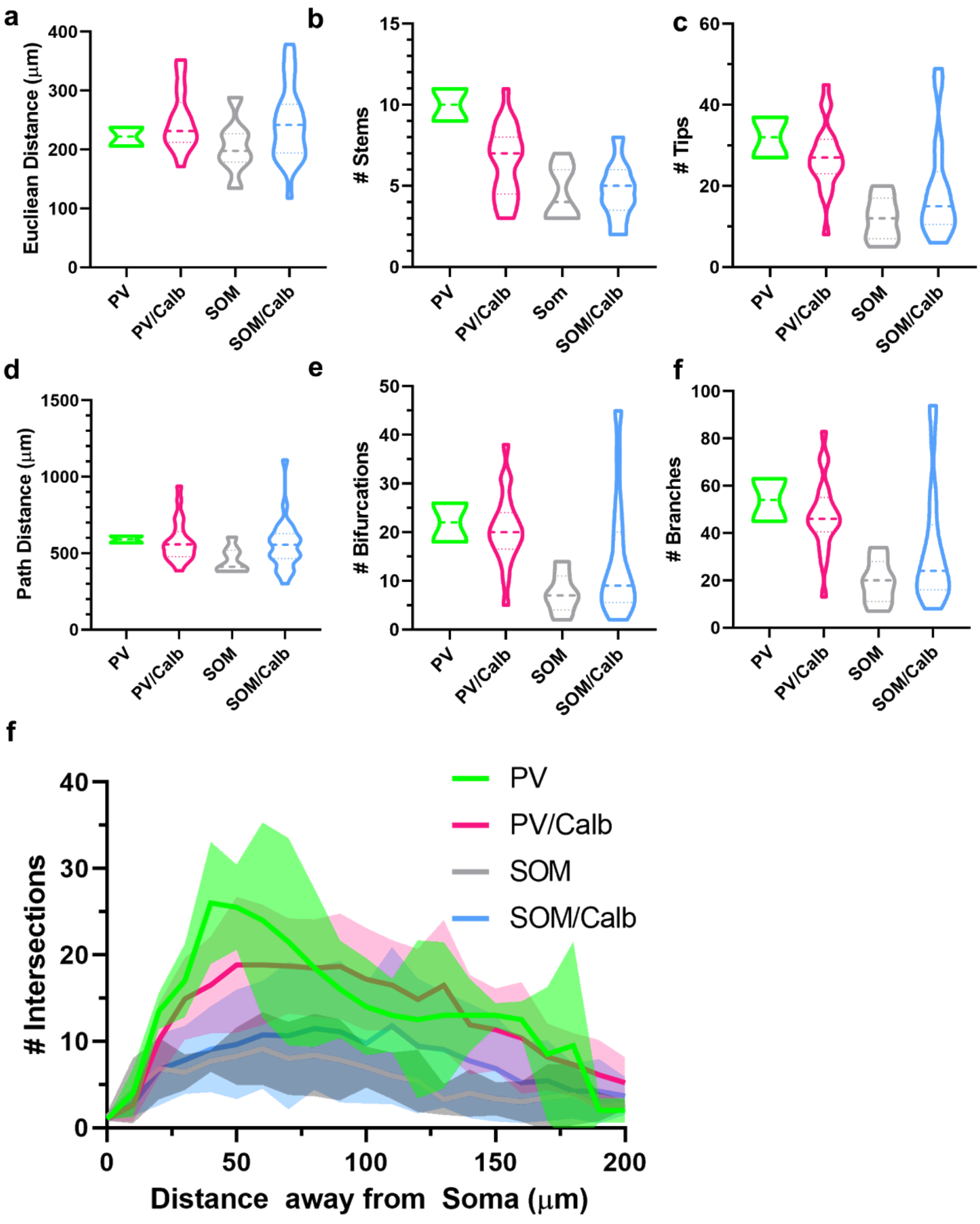


**Supplemental Fig 6. Morphology quantification of 53 inhibitory neurons traced in the BLA.** (**a-f**) Various morphometric parameters were calculated using Vaa3D global neuron feature for each of the 4 molecular subtypes. (**g**) 3D-Sholl analysis for each of the 4 molecular subtypes.

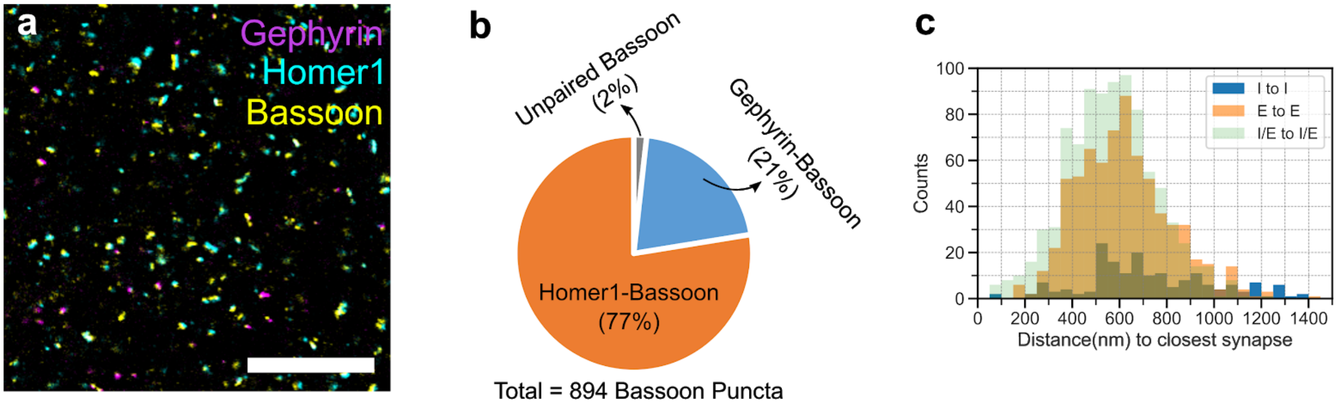


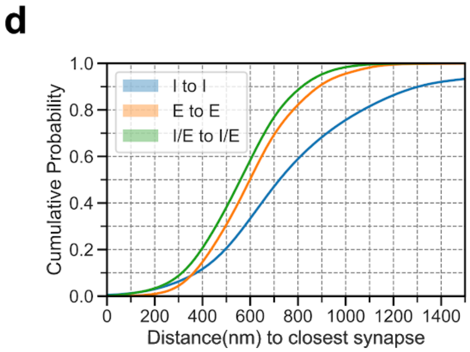


e

| | |
|---|---|
| Gephyrin-Bassoon Pairs/Total Gephyrin Puncta | 1 |
| Homer-Bassoon Pairs/Total Homer Puncta | 0.99 |
| Paired Bassoon/Total Bassoon Puncta | 0.98 |
| Unpaired Bassoon/Total Bassoon Puncta | 0.02 |
| Gephyrin-Bassoon Pairs/Total Bassoon Puncta | 0.21 |
| Homer-Bassoon Pairs/Total Bassoon Puncta | 0.77 |
| Gephyrin-Bassoon Density | 0.27/μm$^3$ |
| Homer-Bassoon Density | 1.01/μm$^3$ |
| Total Synapse Density | 1.28/μm$^3$ |

**Supplemental Fig 7. Homer1 and Gephyrin can be paired with Bassoon immunostaining to represent the majority of excitatory and inhibitory synapses.** (**a**) 100 μm somatosensory cortex tissue was processed with miriEx and triple immunostainied for Gephyrin, Homer1, and Bassoon. The sample was expanded ~4x and imaged. (**b**) All the synaptic pairs were manually annotated within a 15 x 15 x 3 μm$^3$ volume, yielding 706 excitatory synapses (Homer1-Bassoon) and 188 inhibitory synapses (Gephyrin-Bassoon). Pie chart shows that the majority of Bassoon puncta are paired with either Gephyrin or Homer1 in a mutually exclusive manner. (**c**) The distance between inhibitory (I) and excitatory (E) synapses to their nearest neighbor is plotted. (**d**) Cumulative probability plot of the histogram shown in **c**. (**e**) Table of other ratio and density measurements. Scale bars: 5 nm (pre-expansion size).

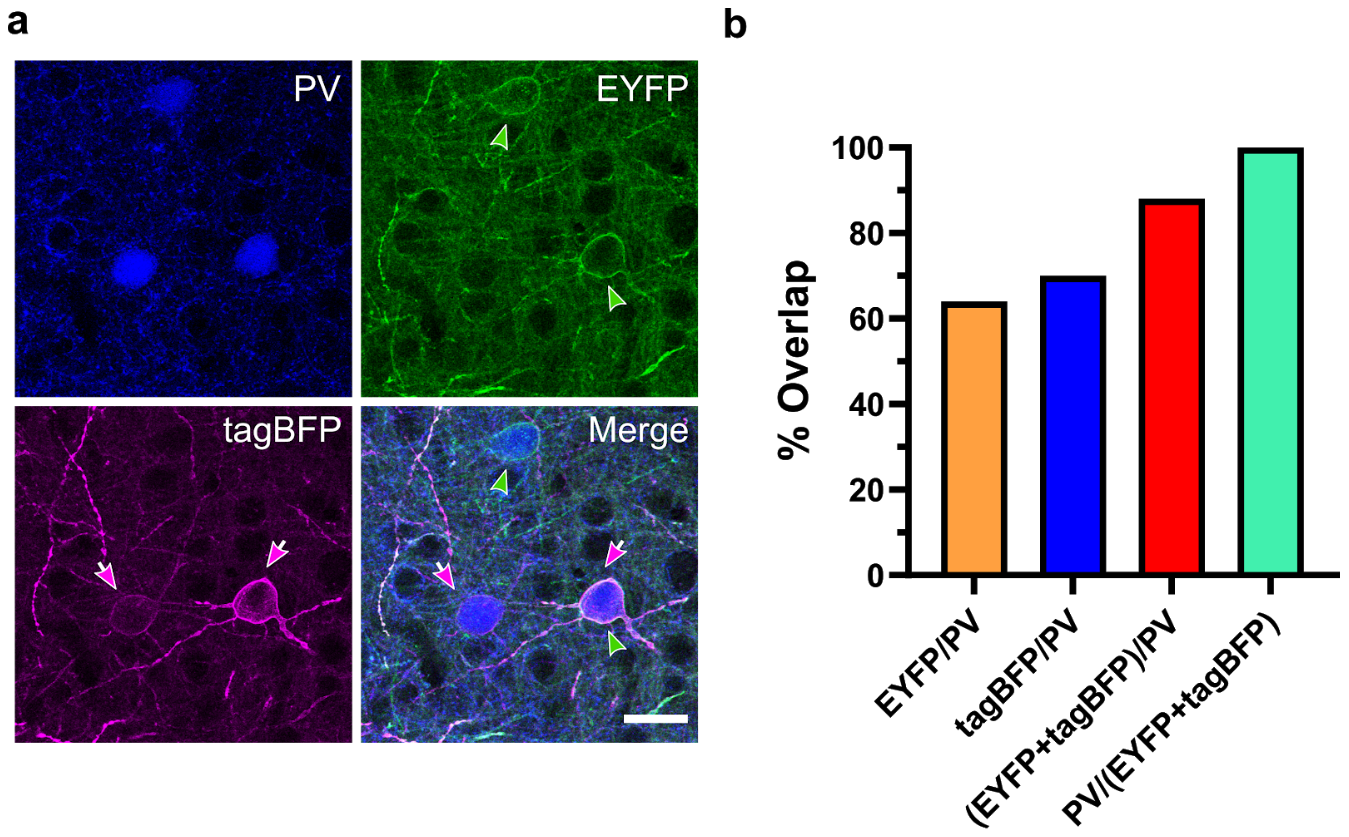


**Supplemental Fig 8. AAV-PHP.eB Brainbow expression in PV-Cre mice is highly sensitive and specific.** (**a**) 100 µm somatosensory cortex tissue was processed with miriEx and triple immunostainied for PV, EYFP, and tagBFP. (**b**) PV neurons across all 6 layers of somatosensory cortex were analyzed for co-expression of PV and 2 out of 4 Brainbow FPs. 88% of all PV neurons (n=50, identified through immunostaining) were positive for either EYFP or tagBFP. 100% of all Brainbow labeled neurons (n=44) were positive for PV immunostaining. PV, parvalbumin; FP, fluorescent protein. Scale bars: 50 µm (pre-expansion size).

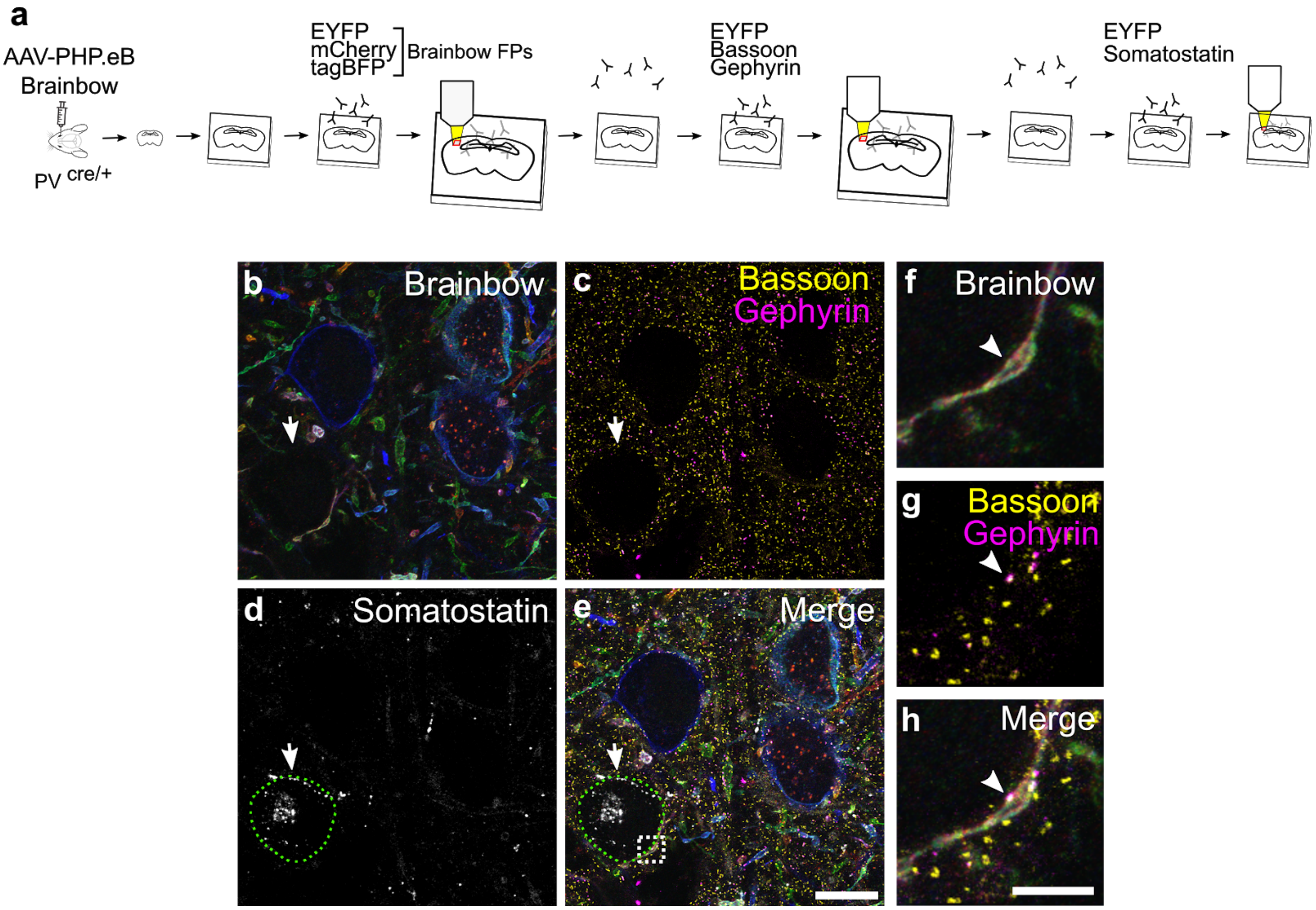


**Supplemental Fig 9. Brainbow FPs, synaptic machinery, and molecular markers can be interrogated in the same piece of tissue.** (**a**) Experimental design: Brainbow FPs, endogenous synaptic markers (Bassoon, Gephyrin), and cell type markers (SOM) are imaged across three rounds of immunostaining using the EYFP channel for registration. (**b**) ~750 nm MIP showing the Brainbow channels imaged in r1. (**c**) ~750 nm MIP showing the Bassoon and Gephyrin puncta imaged in r2. (**d**) ~750 nm MIP showing a Somatostatin positive soma. (**e**) Merged MIP of all three rounds. White arrow points to an "empty hole" that is later defined as a SOM positive soma outlined in a dashed green circle. (**f-h**) Single slice zoomed inset of white square shown in **e**; the arrow points to a putative synaptic connection between the PV axon and soma of the SOM neuron. FP, fluorescent protein; MIP, maximum intensity projection; SOM, somatostatin. Scale bars: (**e**) 10 µm (pre-expansion size). (**h**) 3 µm (pre-expansion size).

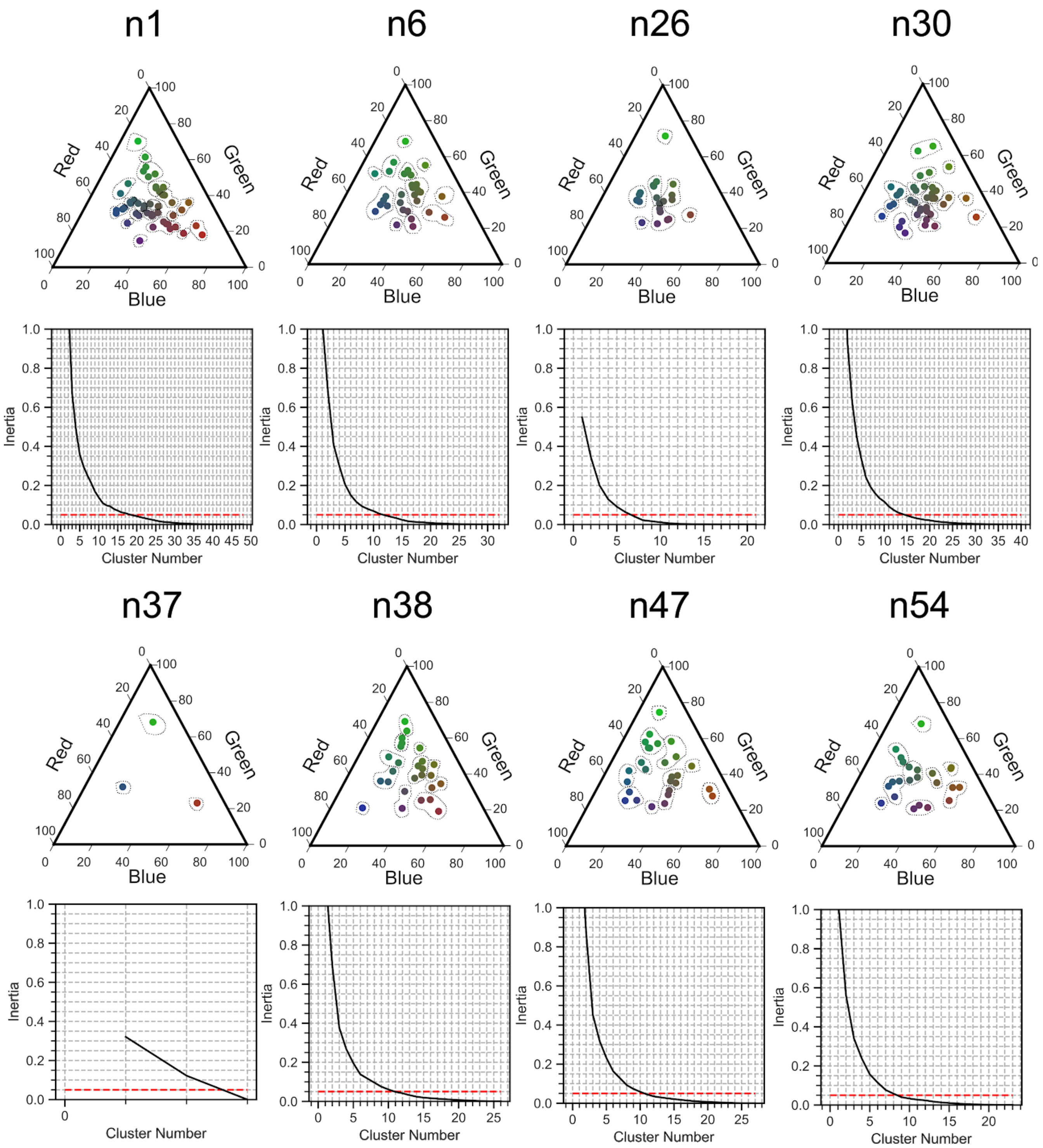


**Supplemental Fig 10. Number of unique axon colors can be identified for k-means clustering.** For each neuron, axon color identities were extracted through nTracer and plotted on a RGB color plot. An elbow plot was used to find the optimal number of k for k-means clustering. This value was conservatively chosen to be where inertia<0.05. The dashed lines represent the k-means color cluster assignments.

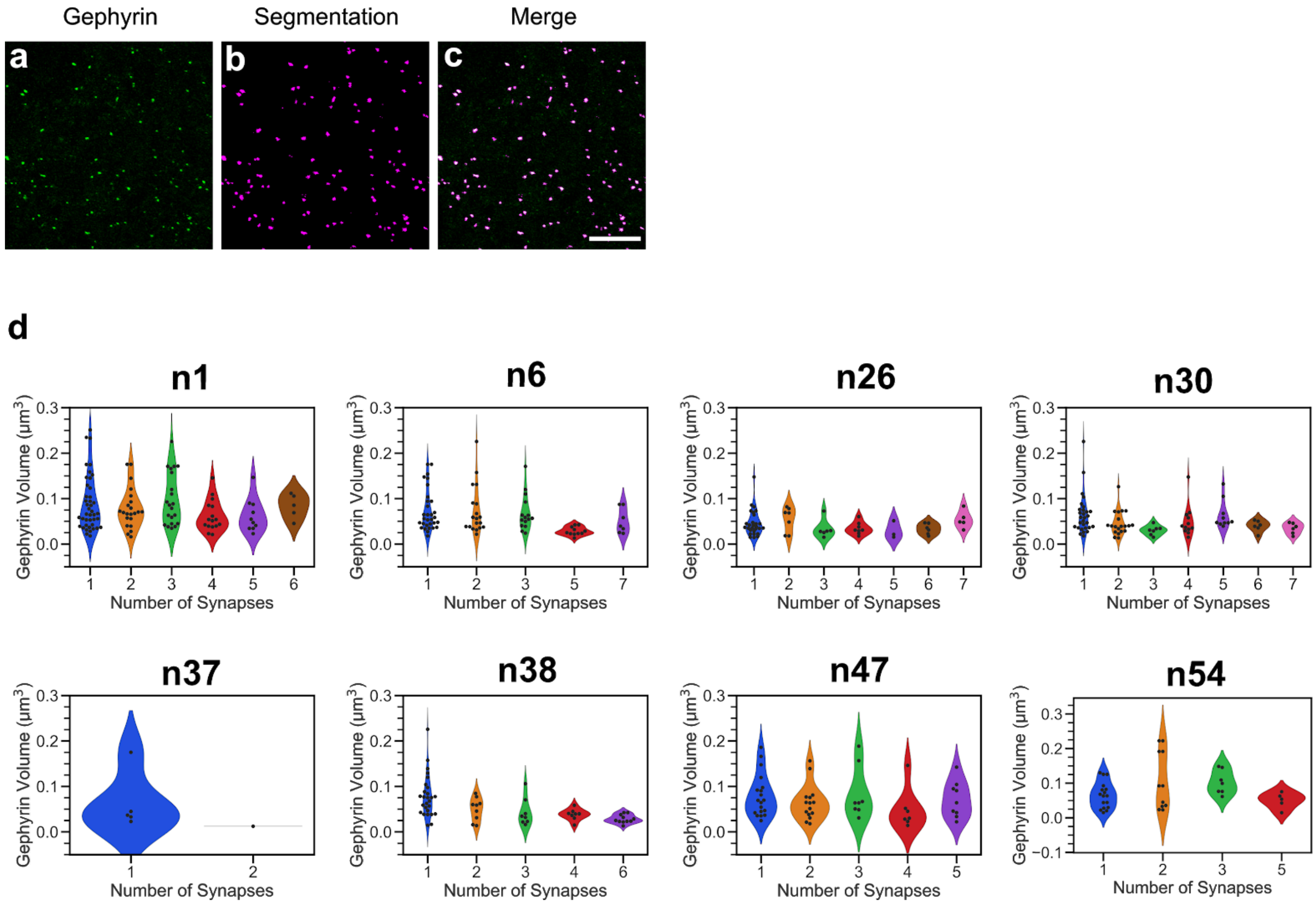


**Supplemental Fig 11. Inhibitory PSD size can be automatically segmented and analyzed.** (**a**) Raw Gephyrin channel. (**b**) Automatic segmentation result. (**c**) Composite image. (**d**) For each individual postsynaptic PV neuron, the size of the PSD was plotted as a function of the number of synapses the presynaptic PV axon formed. Scale bars: 125 nm (pre-expansion size).
