## Supplemental Table 1 for "Light microscopy based approach for mapping connectivity with molecular specificity"

| Figure | Sample | [Aax] | Primary Ab | Primary incubation time and temperature | Secondary Ab | Secondary incubation time and temperature | Imaging | Processing: stitching, chromatic aberration, histogram matching |
|---|---|---|---|---|---|---|---|---|
| Fig 1 | 100 µm BLA from WT mouse | 5 mM | r1: rb-PV (1:500)<br>r2: rb-Calbindin (1:500)<br>r3: rb-CB1R (1:333)<br>r4: rb-PV (1:500)<br>r5: rb-NOS (1:500)<br>r6: rb-SERT (1:500)<br>r7: rb-PV (1:500) | r1-r7: 1d at 37c | r1-r7: rb-AF488 (1:500), DAPI | r1-r7: 1d at 37c | r1-r7: Confocal, 10x Objective (0.4 NA), imaged in 1xPBS | Yes (3x2 tiles), No, No |
| Fig 2 | 200 µm BLA section from PV-Cre/Som-Cre stereotaxically injected with AAV2/9 Brainbow in BLA | 1 mM | r1: rb-Calbindin (1:500), rat-SOMsc(1:100), gp-PV (1:500)<br>r2: rb-mCherry (1:500), sh-GFP (1:500), gp-tagBFP (1:500) | r1: 3d at RT<br>r2: 2d at 37c | r1: rat-AF488(1:500), rb-AF647 (1:500), gp-Cy3 (1:500)<br>r2: rat-AF488(1:500), rb-AF647 (1:500), gp-Cy3 (1:500) | r1: 2d at RT<br>r2: 2d at 37c | r1: Confocal, 10x objective (0.4 NA), imaged in 1x PBS<br>r2: Confocal, 20x Objective (1.0 NA), imaged in 1x PBS | Yes (3x2 tiles), Yes, Yes |
| Fig 3, sFig9 | 100 µm S1 Cortex section from PV-Cre retro-orbitally injected with PhP.eB Brainbow | 1 mM | r1: rb-mCherry (1:500), sh-GFP(1:500), gp-tagBFP (1:500)<br>r2: sh-GFP (1:500), gp-Bassoon (1:500), ms-GephyrinSC (1:100)<br>r3: rat-SOMsc (1:100), sh-GFP (1:500) | r1: 2d at RT<br>r2: 2d at 37c<br>r3: 2d at 37c | r1: rb-AF488 (1:500), sh-Cy3 (1:500), gp-AF647 (1:500)<br>r2: ms-AF488 (1:500), sh-Cy3 (1:500), gp-AF647 (1:500)<br>r3: sh-Cy3 (1:500), rat-AF647 (1:500) | r1: 1d at RT<br>r2: 1d at 37c<br>r3: 1d at 37c | r1: confocal, 20x objective (1.0 NA), imaged in 0.01x PBS<br>r2: confocal, 20x objective (1.0 NA), imaged in 0.001x PBS<br>r3: confocal, 20x objective (1.0 NA), imaged in 1x PBS | No, Yes, Yes |
| Fig 5 | 100 µm S1 cortex section from PV-Cre stereotaxically injected with AAV2/9 Brainbow in BLA | 1 mM | r1: rb-mCherry (1:500), sh-GFP (1:500), gp-tagBFP (1:500)<br>r2: gp-Bassoon (1:500), ms-GephyrinSC (1:100), sh-GFP (1:500) | r1: 2d at 37c<br>r2: 1d at 37c | r1: gp-AF647 (1:500), ms-AF488 (1:500), sh-Cy3 (1:500)<br>r2: rb-AF488 (1:500), gp-AF647 (1:500), sh-Cy3 (1:500) | r1-r2: 1d at 37c | r1: Confocal, 20x Objective (1.0 NA), imaged in 0.01x PBS<br>r2: Confocal, 20x Objective (1.0 NA), imaged in 0.0001x PBS | Yes (2x2 tile), Yes, Yes |
| sFig 1 | 100 µm VGAT-Cre x Ai14 S1 cortex | 1 mM | r1: rb-mCherry (1:500) | r1: 1d at 37c | r1: rb-AF488 (1:500)<br>post strip: rb-AF647 (1:500) | r1/post strip: 1d at 37c | Confocal, 10x objective(0.4 NA) , imaged in 1x PBS | No, No, No |
| sFig 2 | 100 µm Thy1-YFP S1 cortex | 1 mM | sh-GFP (1:500) | 1 day at 37c | sh-AF594 (1:500) | 1d at 37c | Confocal, 10x Objective (0.4 NA), imaged in Vectashield, 1xPBS,0.001x PBS | No, No, No |
| sFig 4 | 100 µm human sensory cortex | 5 mM | r1: rb-Calbindin (1:500), gp-Calretinin (1:500), ms-GFAP (1:500)<br>r2: ms-SMI312 (1:333), rb-PV(1:500) | r1-r2: 1d at 37c | r1: rb-Cy3 (1:500), gp-AF488 (1:500), ms-AF647 (1:500), DAPI<br>r2: ms-AF488 (1:500), rb-AF647 (1:500), DAPI | r1-r2: 1d at 37c | r1-r2: Confocal, 10x objective (0.4 NA), imaged in 1xPBS | Yes (2x1 tile), No, No |
| sFig 5 | 100 µm WT dorsolateral striatum | 5mM | r1: gt-ChAT (1:500), rb-CB1R(1:500), ck-TH (1:500)<br>r2: rb-NOS (1:500), sh-NPY(1:500), rat-SOM (1:100)<br>r3: Lectin (1:1000), ms-GFAP (1:500), rb-SERT (1:500)<br>r4: rb-D2R (1:500), rat-D1R(1:500), gp-Iba1(1:300)<br>r5: gp-VGLUT2 (1:500), rb-MBP (1:500) | r1-r5: 1d at 37c | r1: ck-AF488 (1:500), rb-Cy3 (1:500), gt-CF647 (1:500), DAPI<br>r2: sh-AF488 (1:500), rb-Cy3 (1:500), rat-AF647 (1:500), DAPI<br>r3: ms-AF488 (1:500), rb-AF647 (1:500), DAPI<br>r4: gp-AF488 (1:500), rb-Cy3 (1:500), rat-AF647 (1:500), DAPI<br>r5: gp-AF488 (1:500), rb-Cy3 (1:500) | r1-r5: 1d at 37c | r1-r5: Confocal, 10x objective (NA 0.4), imaged in 1x PBS | Yes (2x2 tile), No, No |
| sFig 7 | 100 µm WT mouse | 1 mM | rb-Homer1a (1:500), ms-GephyrinSC (1:100), gp-Bassoon (1:500) | 2d at 37c | ms-AF488 (1:500), rb-Cy3 (1:500), gp-AF647 (1:500) | 1d at 37c | Confocal, 20x objective (1.0 NA), imaged in 0.001x PBS | No, Yes, Yes |
| sFig 8 | 100 µm S1 Cortex section from PV-Cre retro-orbitally injected with PhP.eB Brainbow (same brain as Fig.3) | 1 mM | gp-tagBFP(1:500), rb-PV (1:500), sh-GFP (1:500) | 1d at 37c | gp-AF647 (1:500), sh-Cy3 (1:500), rb-AF488 (1:500) | 1d at 37c | Confocal, 10x objective (0.4 NA), imaged in 1xPBS | Yes (2x2 tile), Yes, Yes |
