## Supplemental Table 2 for "Light microscopy based approach for mapping connectivity with molecular specificity"

| Target | Vendor | Catalog Number | Host Species |
|---|---|---|---|
| VGAT | Synaptic Systems | 131 011 | Ms |
| Calbindin | Synaptic Systems | 214 002 | Rb |
| PV | Synaptic Systems | 195 004 | Gp |
| PV | Abcam | ab32895 | Gt |
| Homer1 | Synaptic Systems | 160 002 | Rb |
| Gephyrin | Synaptic Systems | 147 111 | Ms |
| Bassoon | Synaptic Systems | 141 004 | Gp |
| CamKII | Abcam | ab22609 | Ms |
| TH | Abcam | ab76442 | Ck |
| NOS | Sigma | n2780 | Rb |
| NPY | Millipore | ab1583 | Sh |
| Somatostatin | Millipore | mab354 | Rat |
| VIP | Immunostar | 20077 | Rb |
| Calretinin | Synaptic Systems | 214 104 | Gp |
| SMI-312 | Biolegend | 837904 | Ms |
| SERT | Synaptic Systems | 340 003 | Rb |
| CB1R | Synaptic Systems | 258 003 | Rb |
| D2R | Synaptic Systems | 376 203 | Rb |
| D1R | Sigma | D2944 | Rat |
| GFAP | Dako Agilent | Z033401 | Rb |
| Vglut2 | Synaptic Systems | 135 404 | Gp |
| IBA1 | Synaptic Systems | 234 004 | Gp |
| MBP | Synaptic Systems | 295 002 | Rb |
| Lectin | Vector Labs | DL-1177 | |
| GAD67 | Millipore | mab5406 | Ms |
| SomatostatinSC | Santa Cruz | YC7 | Rat |
| GephyrinSC | Santa Cruz | G6 | Ms |
| PV | Abcam | ab11427 | Rb |
| NeuN | Millipore | mab377 | Ms |
| Homer1 | Synaptic Systems | 160 006 | Ck |
| GFAP | Sigma | G3893 | Ms |
| CTIP2 | Abcam | ab18465 | Rat |
| tagBFP | Cai Lab custom made | NA | Gp |
| mCherry | Cai Lab custom made | NA | Rb |
| GFP | Biorad | 4745-1051 | Sh |
