## Supplemental Table 3 for "Light microscopy based approach for mapping connectivity with molecular specificity"

| Target | Vendor | Catalog Number | Host Species |
|---|---|---|---|
| Rb-AF488 | Jackson Immunoresearch | 711-545-152 | Dk |
| Rb-Cy3 | Jackson Immunoresearch | 711-166-152 | Dk |
| Rb-AF647 | Jackson Immunoresearch | 711-605-152 | Dk |
| Gp-AF488 | Jackson Immunoresearch | 706-545-148 | Dk |
| Gp-Cy3 | Jackson Immunoresearch | 706-166-148 | Dk |
| Gp-AF647 | Jackson Immunoresearch | 706-606-148 | Dk |
| Sh-AF488 | Jackson Immunoresearch | 713-546-147 | Dk |
| Sh-Cy3 | Jackson Immunoresearch | 713-166-147 | Dk |
| Rat-AF488 | Jackson Immunoresearch | 712-545-150 | Dk |
| Rat-AF647 | Jackson Immunoresearch | 712-606-153 | Dk |
| Ck-AF488 | Jackson Immunoresearch | 703-546-155 | Dk |
| Ck-AF647 | Jackson Immunoresearch | 703-606-155 | Dk |
| Ms-AF488 | Jackson Immunoresearch | 715-546-151 | Dk |
| Ms-Cy3 | Jackson Immunoresearch | 715-166-151 | Dk |
| Ms-AF647 | Jackson Immunoresearch | 715-606-151 | Dk |
